# A single-cell atlas of pig gastrulation as a resource for comparative embryology

**DOI:** 10.1101/2023.08.31.555712

**Authors:** Luke Simpson, Andrew Strange, Doris Klisch, Sophie Kraunsoe, Takuya Azami, Daniel Goszczynski, Triet Le, Benjamin Planells, Nadine Holmes, Fei Sang, Sonal Henson, Matthew Loose, Jennifer Nichols, Ramiro Alberio

## Abstract

Early mammalian gastrulation’s cell-fate decisions are poorly understood due to difficulties obtaining non-rodent embryos. The bilaminar disc of pig embryos mirrors humans, making them a useful proxy for studying gastrulation. Here we present a single-cell transcriptomic atlas of pig gastrulation, revealing cell-fate emergence dynamics, as well as conserved and divergent gene programs governing early porcine, primate, and murine development. We highlight heterochronicity in extraembryonic cell-type development, despite the broad conservation of cell-type-specific transcriptional programs. We apply these findings in combination with functional investigations, to outline conserved spatial, molecular, and temporal events during definitive endoderm (DE). We find early FOXA2+/TBXT-embryonic disc cells directly from DE, contrasting later-emerging FOXA2/TBXT+ node/notochord progenitors. Unlike mesoderm, none of these progenitors undergo epithelial-to-mesenchymal transition. DE/Node fate hinges on balanced WNT and hypoblast-derived NODAL, which is extinguished upon DE differentiation. These findings emphasise the interplay between temporal and topological signalling in early fate decisions during gastrulation.

## Main

The blueprint of the mammalian body plan is laid down during gastrulation, a fundamental process of embryonic morphogenesis that ends with the establishment of the three basic germ layers. Gastrulation can be sub-divided into “primary gastrulation” describing early germ-layer formation events prior to the formation of the node, and “secondary gastrulation” encompassing convergent extension, the onset of neurulation and somitogenesis^1^. The unfolding of these processes has been mapped using single-cell transcriptomics in the mouse^2^, rabbit^3, 4^, non-human primates^5, 6^, and partially in humans^5^. Cross-species analyses have identified broadly conserved and divergent features of major lineage emergence, however, detailed investigations of “primary gastrulation” are limited due to the scarcity of cells in these datasets. Furthermore, validation of transcriptomic observations using embryos is limited by the lack of specimens in non-human primates and humans. To address this, here we present a high-resolution single-cell transcriptomic atlas of pig gastrulation and early organogenesis, comprised of 91,232 cells from 62 complete pig embryos collected between embryonic days (E) 11.5 to 15 (equivalent to Carnegie stages (CS) 6 to 10). The pig embryo, like most other mammals, forms a flat embryonic disc (ED) prior to the onset of gastrulation and represents an accessible species for functional investigations^7^. Importantly, the pig is a valuable model for biomedical research and is increasingly being utilized for the development of transplantable organs for humans^8–10^.

We used this comprehensive dataset to shed new light on the salient features of gastrulation in mammals. By performing cross-species comparisons we uncover heterochronic differences in the development of extra-embryonic cell types. Despite variability in differentiation dynamics and pathways regulating cell behaviour, there is broad conservation in cell type-specific programs across pigs, primates and mice. We focussed on the long-standing question of how the definitive endoderm (DE) emerges during gastrulation in mammals. Despite the evidence of mesendodermal progenitors in invertebrates, fish and chick^11–14^, recent studies in the mouse demonstrated that epiblast cells give rise to DE independent of mesoderm^15, 16^. However, evidence from studies using mouse and human embryonic stem cells (hESC) suggests that a common mesendodermal progenitor may also exist in mammals^17–20^. We combined transcriptomic analysis and embryo imaging to show that soon after the first mesodermal cells appear in the posterior epiblast a group of ED disc cells expressing FOXA2+ delaminate to give rise to DE, these cells differ from later FOXA2-TBXT+ cells which give rise to the node/notochord. Further, both cell types form via a mechanism independent of mesoderm and do not undergo an epithelial-to-mesenchymal transition (EMT). Further, functional validations using *in vitro* differentiation of pluripotent pig embryonic disc stem cells (EDSCs) and hESC^21, 22^ show that a balance of WNT and Activin/NODAL signalling are critical to the acquisition of the endoderm fate. Together, our findings indicate that the temporal dynamics and spatial localisation of WNT, originating from the primitive streak, coupled with hypoblast-derived NODAL play critical roles in orchestrating primary gastrulation in mammals.

## Results

### Single-cell transcriptome of pig gastrulation and organogenesis

To investigate cellular diversification during gastrulation and early organogenesis in flat-disc embryos, we obtained scRNA-seq profiles from 23 pooled samples encompassing 62 pig embryos, collected at twelve-hour intervals between E11.5 and E15 using the 10X chromium platform (Fig. 1a; Extended Data Fig. 1). The dataset includes early streak up to 12-somites stage, equivalent to Carnegie stages (CS) 6 to 10 (Fig. 1b). Transcriptomes of 91,232 cells passed quality controls, with a median of 3,221 genes detected per cell (see Methods; Extended Data Fig. 1a-c). known cell-type markers and unbiased clustering of all samples (See Supplementary Table 1) were used to identify 36 major cell populations and concomitantly, novel pig cell-type marker genes (see Methods, Fig. 1c; Extended Data, Fig. 1d-e; Supplementary Table 2). Early embryonic cell types, such as epiblast and primitive streak (PS) cells, decreased in number over time concomitant with their differentiation (Fig. 1d). Mesoderm and DE progenitors were present as early as E11.5 suggesting that gastrulation may commence before the morphological changes due to A-P patterning become visually apparent. Most mesoderm diversification occurs from E12 while ectodermal lineages appear from E13.5 onward (Fig. 1d). Extra-embryonic mesoderm (ExM) cells were identified from the earliest timepoint (E11.5), in agreement with reports that ExM emerges prior to the formation of the PS in primates and pigs^23–25^. Consistent with the well-documented late emergence of amnion in most domestic animals compared to primates, the amnion cluster was not present in earlier samples but appeared later in gastrulation, from E12.5 onward (Fig. 1d). While it is possible that amnion may be involved in cell patterning in pig, as in primates, a role in A-P patterning is unlikely, as this occurs prior to amnion formation^26, 27^.

**Fig. 1.**
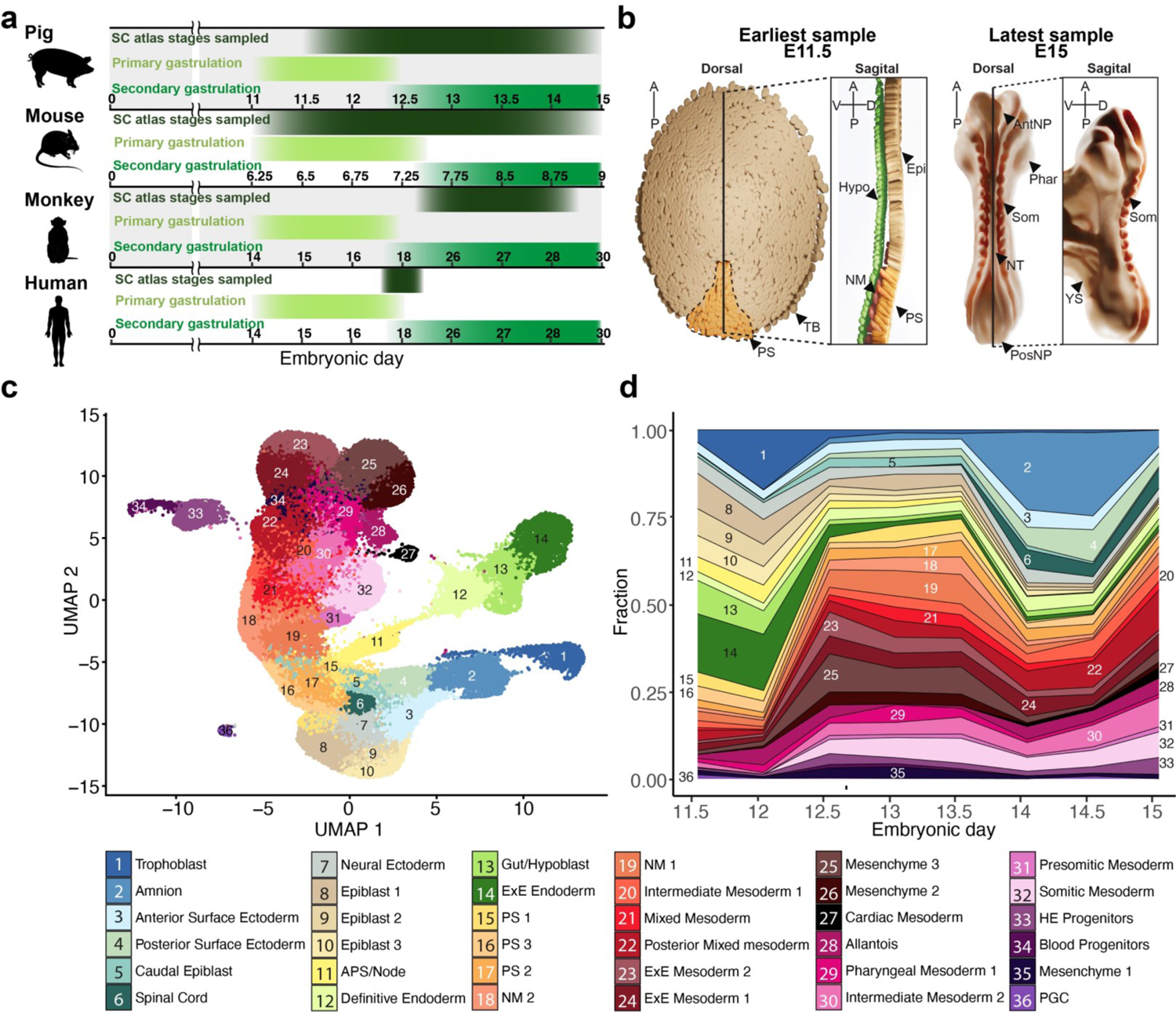
Overview of the pig single cell atlas. **a**, Schematic showing onset and duration of primary and secondary gastrulation in humans, mice, pigs and monkeys. Timepoints where high-resolution single-cell datasets are available, are marked for each species^2,6,28^ as well as the time points covered in this atlas. Numbers indicate embryonic day. **b**, Diagrammatic representations of the earliest and latest embryo samples in this dataset with visible embryonic structures/cell types labelled. Epi, Epiblast; PS, Primitive streak; NM, Nascent mesoderm; Hyp, Hypoblast; TB, Trophoblast; NT, Neural tube; Phar, Pharyngeal arches; Som, Somites; PosNP, Posterior neuropore; AntNP; Anterior neuropore; YS, yolk sac. **c**, Uniform manifold approximation and projection (UMAP) plot showing atlas cells (91,232 cells). Cells are coloured by their cell-type annotation and numbered according to the same legend as **d** below. **d**, Stacked area plot showing the fraction of each cell type at each time point, a progressive increase in cell-type complexity can be seen across time points with mesodermal cell type diversification preceding that of ectoderm. APS, Anterior Primitive Streak; ExE, extra-embryonic; PS, Primitive streak; NM, Nascent mesoderm; HE, Hematoendothelial; PGC, Primordial germ cells.

### Pig embryos show broad similarities to human, monkey and mouse embryos

To gain insights into conserved and divergent features of non-rodent and rodent mammals we compared the transcriptomes of peri-gastrulation stage mouse^2^ and *Cynomologous* monkey^6^ embryos with pigs using high-confidence one-to-one orthologues (see methods). Label projection of our pig dataset onto mouse^2^, showed that the majority of cell-type annotations were well-matched between both species (Fig. 2a), with large fractions of each cell type allocated as being analogous to their mouse counterparts including cardiac mesoderm, mesenchyme 2&3, extraembryonic endoderm, spinal cord, primordial germ cells (PGCs), and epiblast 2 (Extended Data Fig. 2a). By contrast, fewer cells from transient mesodermal progenitors such as nascent, posterior mixed mesoderm and intermediate mesoderm were allocated an analogous identity. Given that these progenitors are generally transcriptionally similar and identified by broad markers common in multiple mesodermal progenitors (e.g. *MESP1, HAND1* or *OSR1*) this discrepancy may be a result of these cells having few specific markers. In the case of extra-embryonic tissues, such as amnion and trophoblast, a large fraction of these tissues was allocated surface ectoderm identity. Similarly, a large portion of ExM was allocated as mesenchyme. Projections of pig stages onto mouse development show our time course closely aligns between E7.25 to E8.5 (Fig. 2b). Projection mapping to the macaque dataset also showed a high degree of similarity of cell type annotations; in contrast to mouse-annotation mapping, this resulted in better agreement between predicted identities and our own annotations of extra-embryonic mesodermal tissues (Fig. 2c-d; Extended Data Fig. 3). Given the large discrepancies in the methodologies and criteria used for annotating cell types in single-cell data^28^ meaningful cross-species comparisons can often be challenging. To overcome these challenges, we used data projection/label transfer to apply consistent annotation to equivalent cell types for further cross-species comparisons.

**Fig. 2.**
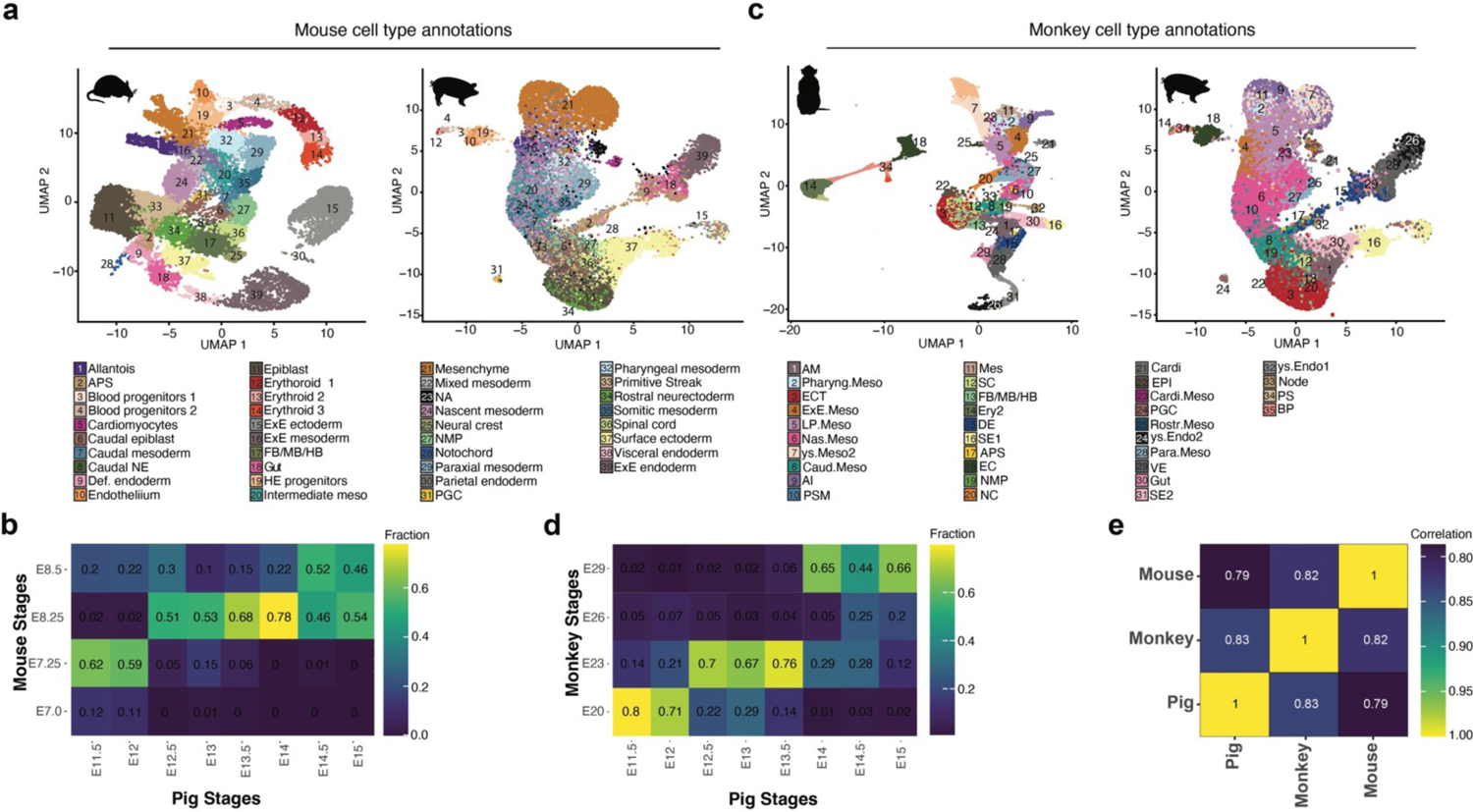
Alignment of Pig, Mouse and Monkey datasets. **a**, UMAPs showing E6.5–8.5 mouse embryo cell types (left)^2^ and Pig E11.5 to E15 with mouse annotations (right) after reciprocal PCA-based projection onto the mouse dataset. **b**, Heat map showing the fraction of pig cells by stage allocated to a particular mouse stage after label transfer. **c**, UMAPs showing E20-29 monkey embryo cell types (left)^6^ and Pig E11.5 to E15 with mouse annotations (right) after reciprocal PCA-based projection onto the monkey dataset. **d**, Heat map showing the fraction of pig cells by stage allocated to a particular monkey stage after label transfer. **e**, Heatmap showing the Pearson correlation co-efficient between the transcriptomes of pig, monkey and mouse embryos.

We calculated correlation co-coefficients between the transcriptomes of monkey, mouse and pig embryos and showed that generally both pigs and mice correlated more closely to monkeys than to each other (Fig. 2e). Hierarchal clustering of individual cell types generally grouped cell types with lower correlation together, these included several extra-embryonic tissues (e.g. ExE Endo 1, 2, 4, 5 and Hypoblast 1 and 2) together corroborating known differences in the morphology and regulation of these tissues^3, 27, 29^. We noted that among cell-type specific marker genes, there was a large degree of overlap between monkeys, mice and pigs. This allowed us to determine sets of highly conserved cell type-specific markers: epiblast 1 (*POU5F1, SALL2, OTX2, PHC1, FST, CDH1 and EPCAM*), PS 1 (*CDX1, HOXA1, SFRP2, and GBX2*), APS (*CHRD, FOXA2, GSC, CER1 and EOMES*), node (*FOXA2, CHRD, SHH and LMX1A*) (Extended Data Fig. 4), DE/Foregut (*SOX17, FOXA2, PRDM1, OTX2 and BMP7*) and DE/Hindgut (*SOX17, FOXA2, TNNC1 and ITGA6)*. Notably, we also identified sets of genes that were strong cell type identifiers in monkey and pig cell types, but not mice, for example, *UPP1, SFRP1, PRKAR2B, APOE and IRX2* in the epiblast, *CD9, GPC4* and *COX6B2* in the APS and *PTN, HIPK2* and *FGF8* demarcating Node. We identified conserved and divergent markers for other less well-characterised cell types (Supplementary Tables 3-5). These observations highlight the importance of investigating multiple representative animal models to identify conserved gene-regulatory networks relevant to cell fate decisions in mammals.

We then looked for transcriptional differences outside of cell-type specific gene programs, utilising ClusterProfiler to elucidate KEGG term enrichment among differentially expressed genes (DEGs). This revealed a considerable number of genes that were markedly upregulated in pig and monkey epiblasts compared to mice (Extended Data Fig 5a). Further examination showed that a significant proportion of these DEGs were replicated across multiple cell-type comparisons. Notably, many of the identified genes were associated with distinct signalling pathways, including the Mitogen-Activated Protein Kinases (MAPK) and the Phosphatidylinositol 3-Kinases (PI3K)/Akt pathways, along with cell adhesion pathways such as those mediating focal adhesions and adherens junctions (Extended Data Fig 5b). Given that these differentially expressed genes are part of pathways generally implicated in cell behaviour such as the regulation of cell growth, proliferation, differentiation, and morphogenesis, this may correspond to known differences in embryo size, cell cycle length and morphology between these species.

We next used our dataset to better understand the process of human gastrulation, currently informed by a single gastrula-stage CS7 (E17-19) human embryo^30^. Cross-species reference mapping of the CS7 human embryo onto our own dataset as well as that of mouse and monkey revealed distinct asynchronicity between cell differentiation dynamics across species. Intriguingly, despite the relatively immature stage of the human embryo, the mapping of mesodermal cell types onto their porcine counterparts revealed a considerable degree of alignment with E15 extra-embryonic mesoderm. This alignment potentially suggests that human extraembryonic mesoderm not only envelops the embryo more extensively but also exhibits accelerated maturation when compared with other cell types, such as the epiblast and nascent mesoderm clusters. The latter two were found to correspond more closely with their E13 porcine equivalents, which more closely mirror the morphology of a CS7 human embryo. A congruous trend was observed comparing the human embryo to mice^2^, as the human yolk sac mesoderm aligned closely to E8.5 mesenchyme (Fig. 3a). Comparisons of endodermal cell types also showed asynchronous development of yolk sac endoderm, like ExE mesoderm, human yolk sac endoderm had a higher mapping frequency to pig E15 yolk sac endoderm and E8.5 ExE endoderm (Fig.3b). By contrast, nearly all the cell types investigated showed that CS7 human cell types matched CS9 in primates (Extended Data Fig. 6a). While these results might reflect a discrepancy in embryonic staging, they also suggest fewer asynchronicities between human and monkey embryos. We did, however, note that ectodermal tissues such as amnion and surface ectoderm appeared to differ between human and monkey annotations, while the pig annotations of these tissues aligned more closely to human suggesting there is a need to better define the transcriptional profiles of these tissues for further comparisons. Together these findings have a key implication, namely that while the cell-type specific differentiation programs themselves show a remarkable amount of conservation differentiation dynamics appear to differ across species.

**Fig. 3.**
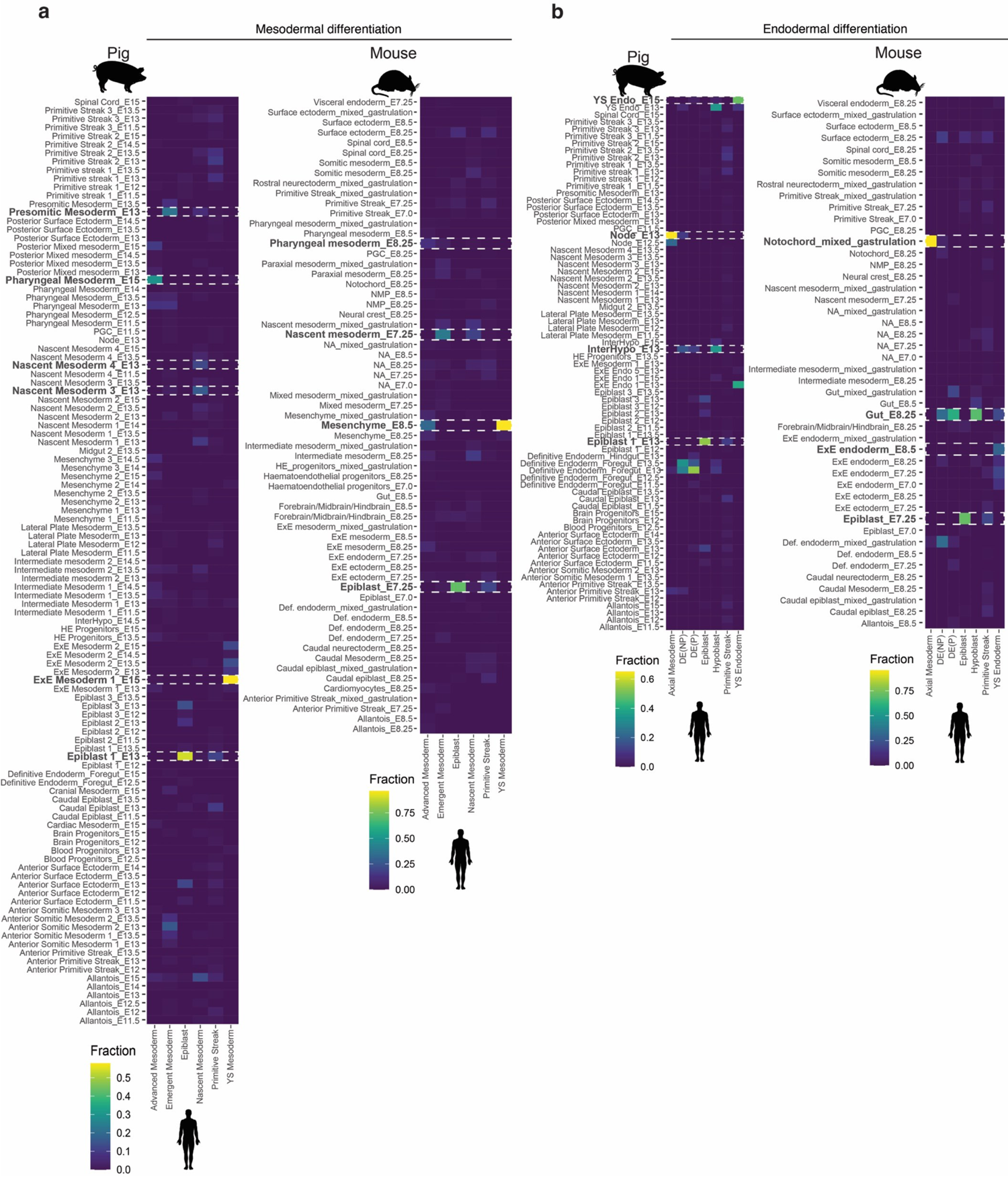
Heterochrony across Pig, Mouse and Monkey differentiation. **a**, Heat maps showing the fraction of human mesodermal cells^30^ grouped by cell type allocated to a pig (left) or mouse^2^ (right) cell identity after label transfer. **b,** As with a, except with endodermal cell types. A fraction of 1 indicated that 100% of cells of a given cell type were predicted to be analogous to the cell identity in the queried organism.

### Mesodermal and endodermal derivatives of the porcine epiblast are transcriptionally distinct

Given that our results suggested conservation in cell-type specific programs, we reasoned that the core mechanisms of differentiation would be conserved between mice and large mammals. One area of controversy is that of endodermal differentiation. Indeed, it has recently been suggested proliferative, bi-fated “mesendodermal” progenitors have not been identified in the mouse embryo^15, 16, 31^. However, this idea has gained less traction in large mammalian embryology as early *in vitro* evidence in hESCs has suggested that bipotent progenitors may exist^17–20^. Given the high numbers of cells from early gastrulation, our dataset allowed us to dissect the events of this period at high resolution. We analysed epiblast, PS, APS/node, nascent mesoderm, and DE clusters (Fig. 4a-d). Sub-clustering of mesoderm and endoderm-fated cells identified 11 subpopulations: four nascent mesoderm, three PS, APS, node, DE, and early gut (Fig. 4b). Nascent mesodermal cells expressing *MESP1* continued to increase in number between E11.5 to E15, while PS and APS clusters decreased (Fig. 4c-d). High-*SOX17* DE cells were stable across the time course. In contrast, the APS-neighbouring node cells predominantly emerged one day later (E12.5). This finding is in line with previous observations where the *GSC*-expressing node can be identified from E12-E13 pigs^32^. Of the PS clusters, PS1, which corresponds to pre-streak posterior epiblast, showed higher *EOMES* expression than PS2 and PS3. PS2 and PS3 had markedly higher expression of *CDX1* and *HAND1* respectively, suggesting that PS2 and PS3 represent mesoderm progenitors. We observed that most cells within the APS expressed both *FOXA2* and *CHRD*. Notably, however, the APS cluster manifested certain heterogeneity. A significant proportion of cells exhibited elevated expression levels of markers such as *POU5F1, NANOG, EOMES,* and *CER1*. Conversely, a subset of cells displayed diminished levels of these same markers, coupled with heightened expression of TBXT. SCENIC analysis also confirmed that cell subtypes demonstrated unique regulon enrichment profiles (Fig. 4e). These observations are consistent with the idea that distinct populations of the APS may give rise to the DE and node.

**Fig. 4.**
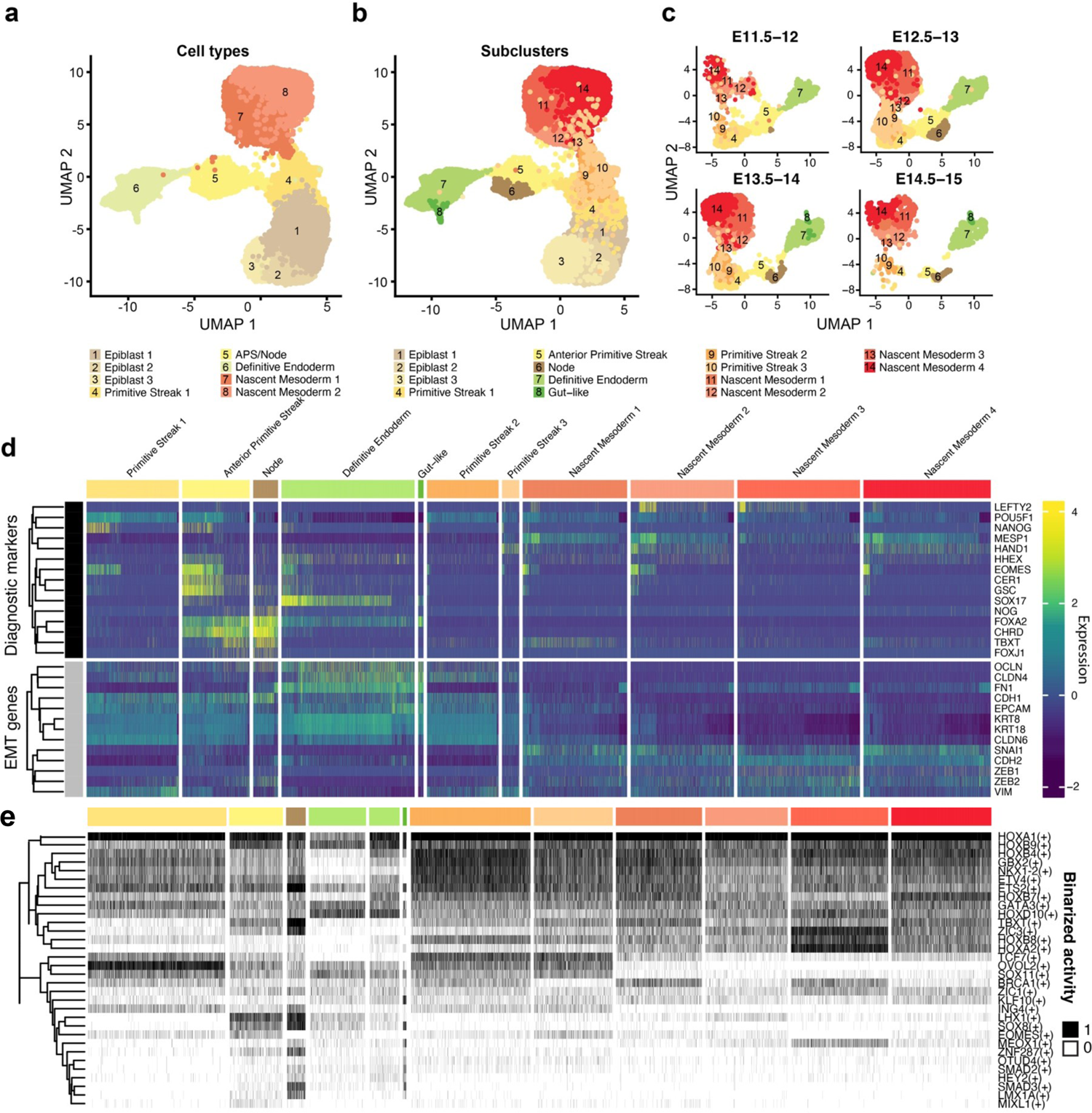
Endodermal progenitors do not undergo EMT. **a**, UMAP plot showing Epiblast, PS, APS/Node, DE and nascent mesoderm clusters (11,450 cells; E11.5-E15) coloured by global cell-type annotation. **b**, As with a, coloured by cell subtypes. **c**, UMAP plots showing identified subclusters at specific time points in development. E, Embryonic day. **d**, Heat map illustrating the scaled mean expression of genes. Expression of selected markers used to identify cell subclusters (top) as well as epithelial and mesenchymal marker genes (bottom). **e**, Binarized heatmap showing SCENIC regulon activity in individual cells. Cell type legend is the same as d.

We next analysed the expression of epithelial markers and genes related to EMT (Fig. 4d). Tight junction markers, such as *OCLN, CLDN6,* and *CLDN4*, along with the intermediate filament protein-encoding genes *KRT8* and *KRT18*, displayed lower expression within nascent mesodermal clusters. Except for *CLDN6*, these markers also exhibited higher expression in the DE cluster. As with other epithelial markers, *CDH1* and cell-cell adhesion-associated *EPCAM* also showed diminished expression within nascent mesoderm. In contrast, the expression of the mesenchymal transition regulator *SNAI1* showed elevated expression within the nascent mesodermal clusters. Intermediate filament and mesenchyme marker *VIM* displayed parallel expression profiles in both nascent mesoderm and pre-streak (PS) clusters, including PS1. Unexpectedly, extracellular matrix protein-encoding *FN1*, frequently associated with EMT, exhibited substantial expression within the DE. Likewise, *ZEB2*, a transcriptional repressor of *CDH1*, was expressed in both nascent mesodermal clusters and the node.

Together these data suggest that cells of the posterior epiblast (PS1), along with the APS, node, and DE, retain a robust epithelial identity throughout their differentiation process. This epithelial-to-epithelial transition has been previously described during the formation of the amnion in the mouse^33^. In contrast, nascent mesodermal undergoes a divergent process resembling the “classic” model of EMT. Therefore, the processes by which epiblast cells transition from a columnar to a simple epithelium (in the case of DE), or toward a mesenchyme/mesenchyme-like state, as is the case for nascent mesoderm and notochordal process respectively, appear to be highly nuanced and tissue-specific. This observation is especially pertinent for the DE and nascent mesoderm, as we found no evidence supporting a common mechanism of cell ingression, in line with findings in mice^15, 16, 34^.

### Exploring early somitogenesis in pig embryos

To explore the derivatives of the nascent mesoderm cells we sub-clustered nascent mesoderm, pre-somitic and somitic mesodermal cell types (Extended Data Fig. 7). This facilitated the identification of seven subtypes: 3 anterior somitic mesoderm clusters, cranial mesoderm, dermomyotome/sclerotome, posterior somitic mesoderm and pre-somitic mesoderm. We observed several genes with similar dynamics in pigs, as reported in mice and in human *in vitro* models. For example, *TBX6*, is highly expressed in pre-somitic mesoderm and posterior somites, but less so in more mature somitic cell types^35–37^. In contrast, *MYF5* and *MYOD*, were expressed at later time points. Additionally, *FOXC2* was lowly expressed in all somitic subtypes. Generally, we observed the first mature somite subclusters emerge from day 14 onward while presomitic mesoderm clusters were present throughout, consistent with patterns described in many other species. We have also noted several new markers of pig somitogenesis (Extended Data Fig. 7d).

### Identification of FOXA2 and TBXT cells in gastrulating pig embryos

To spatially position mesodermal and endodermal progenitors, we conducted whole-mount immuno-fluorescence imaging of E10.5-E11.5 porcine embryos for SOX2, FOXA2, and TBXT (Fig. 5; Supplementary Videos 1-3. At E10.5, TBXT-positive (T+) cells appear in the posterior end of the epiblast, their number are increased in E11.5 embryos. Many of these cells extend beyond the posterior ED boundary as ExM by the end of E11.5 (Fig. 5a-d). A group of FOXA2*-*positive (F+) cells was detected anterior to the FOXA2+ and TBXT+ (FT+) cells at E11.5. A similar population of F+ cells in the epiblast layer was recently reported in mice^15, 16^. While initially this F+ population outnumbered FT+ cells by a factor of ∼3, this gradually decreased to ∼1.4 by the end of E11.5 (Fig. 5c-d). Lateral views of the reconstructions showed F+ cells part-way delaminating from the epiblast into the hypoblast layer (Fig. 5e). Conversely, we did not observe any FT+ cells intercalating. In line with their spatial positioning and the transcriptome analysis (Fig. 4) these observations suggested the F+ population is fated to DE while the FT+ cells may represent node/notochord precursors. This is further supported by the increased number of double-positive TBXT/FOXA2 (FT+) cells demarcating the medial APS region by the end of E11.5.

**Fig. 5.**
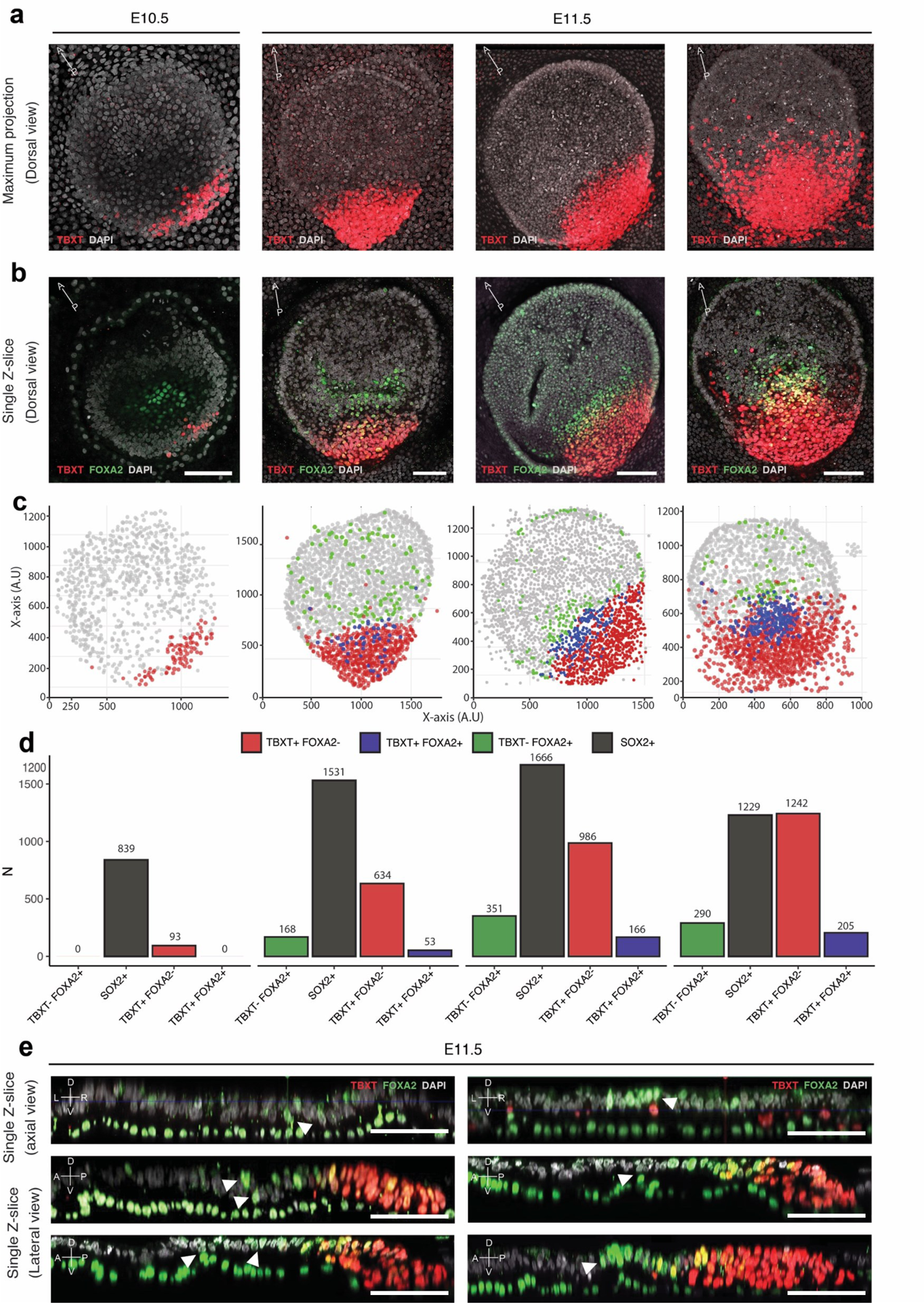
FOXA2 and TBXT domains are spatially separated. **a**, Dorsal view maximum projection of E10.5 to E11.5 porcine embryos showing TBXT expression. E11.5 embryos are ordered youngest to oldest from left to right. **b**, Single z-slice of the embryos shown in a, showing FOXA2 and TBXT expression. **c**, *In Silico* representations of embryos following 3D segmentation of embryos from a and b. **d**, Quantification of TBXT, FOXA2, and SOX2 expressing cells in embryos from a and b. **e**, Axial and lateral reconstructed sections of embryos stained for FOXA2 and TBXT. White arrowheads indicate FOXA2+ TBXT-cells that are spatially separated from the TBXT domain. Scale bars: 100µm

### Spatiotemporal events and transcriptional signatures identify mesodermal and endodermal during primary gastrulation

To investigate whether spatially separated F+ and FT+ cells acquired alternative fates, we isolated posterior epiblast PS1, APS, node and DE clusters and ordered them in pseudo-time (Fig. 6a-b). As expected PS1 and APS cells were ordered at the start of the trajectory before bifurcating toward node or DE fates. To directly infer the fates of the cells observed by IF, we classified cells based on FOXA2 and TBXT expression (Fig. 6c). We previously showed that NANOG and SOX17 are exclusively expressed in the ED and hypoblast layer respectively in pig E10-E11.5 embryos^38^, thus we reasoned that including the expression status of these markers in conjunction with FOXA2 and TBXT would allow us to ascertain the position of cells during fate commitment. We identified, 143 FOXA2+, NANOG+, TBXT-, SOX17-(FN+) cells, 73 FOXA2+, NANOG+, SOX17+, TBXT-(FNS+) and 505 FOXA2+, TBXT+, NANOG-, SOX17-(FT+) cells (Fig. 6c). While we identified 136 FOXA2/TBXT/SOX17 (FTS+) positive, their transcriptional profiles were closely aligned with those of the FT+ cells (Extended Data Fig. 8a, Supplementary Table 6).

**Fig. 6.**
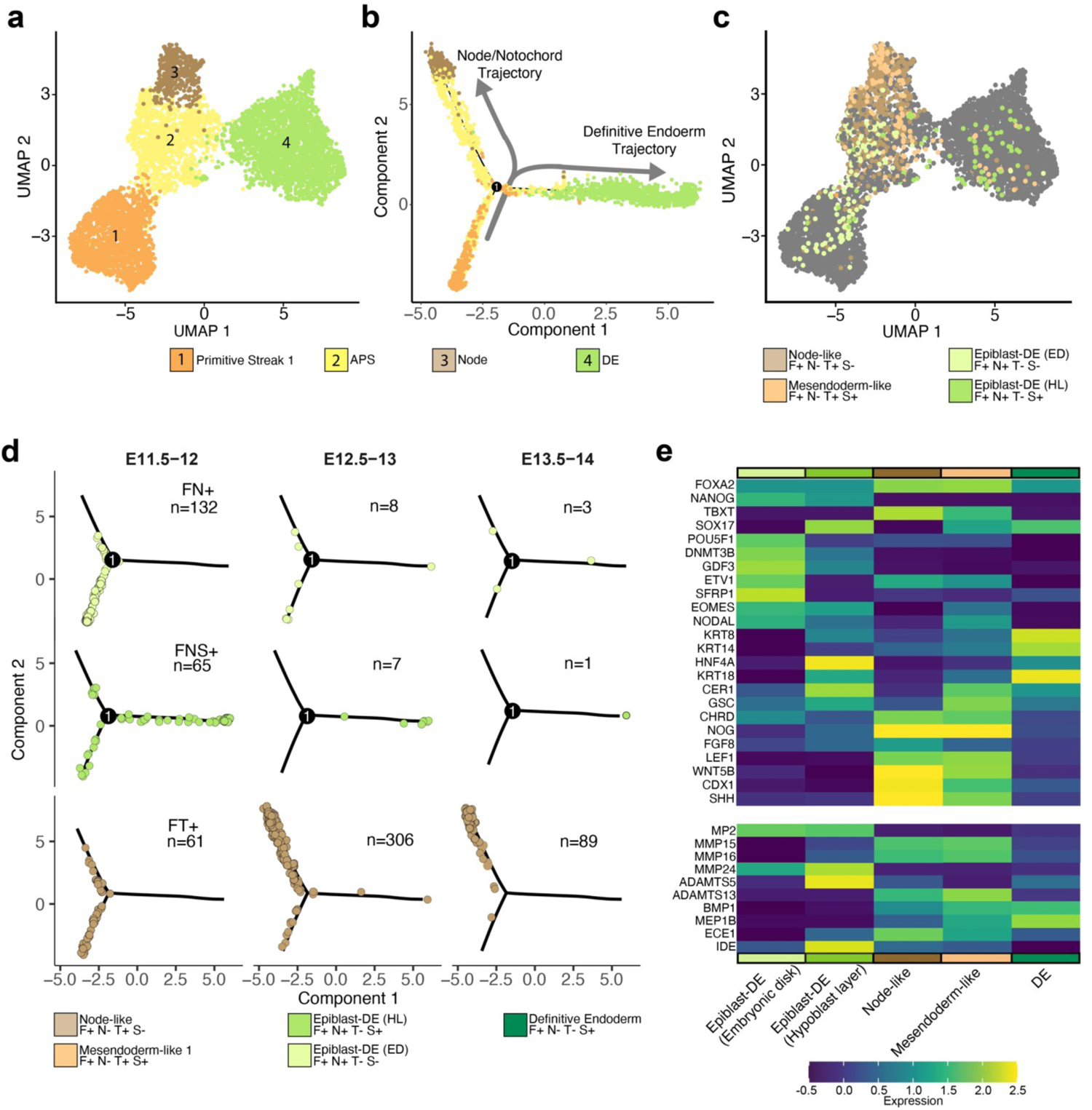
Endoderm forms from Epiblast-like TBXT-progenitors. **a**, UMAP plot showing Primitive Streak 1, Anterior Primitive Streak, Node and Definitive Endoderm subclusters (5,860 cells) coloured by cell-type. **b**, Force-directed graph of the cells illustrated in a, after pseudo-temporal ordering using Monocle2. **c**, UMAP plot categorised by FOXA2, NANOG, TBXT and SOX17 expression. F, FOXA2; N, NANOG; T, TBXT; S, SOX17 cells. Cells are coloured by their F/N/T/S category. **d**, Force-directed graphs of the cells illustrated in c, after pseudo-temporal ordering using Monocle2 and splitting into developmental time point bins. **e**, Heat map illustrating the scaled mean expression of genes. Top: selected cell-state markers from significantly DEGs in the categorically defined cells in c. Bottom: expression of selected metalloproteases.

Pseudo-temporal ordering revealed that FN+ cells represent uncommitted cells prior to the node/DE bifurcation trajectory. FNS+ cells, by contrast, were present both early in the trajectory and throughout the DE trajectory (Fig. 6d). Notably, most FTS+ and FT+ cells from E12.5 onward were biased toward node/notochord. Together this suggests that TBXT-cells primarily contribute to DE. Differentially expressed genes (DEGs) between these groups confirmed the ED-like identity of FN+ cells, as other epiblast markers including *POU5F1, DNMT3B, ETV1, GDF3* and *SFRP1* were significantly (FDR > 0.0001) elevated (Fig. 6e, Supplementary Table 6). Notably, FN+ cells express higher levels of specific matrix metalloproteases (Fig. 6e), suggesting that these enzymes may facilitate their passage through the basal membrane. DEGs in FNS+ cells showed that despite *NANOG* expression, these cells had significantly (FDR > 0.0001) elevated levels of *CER1*, *HNF4A* and epithelial DE markers including *KRT8* and *KRT18.* Albeit these were considerably lower than *NANOG*-DE cells, consistent with the idea that these cells represent the earliest DE cells (Fig. 6e). FTS+ and FT+ cells showed significantly (FDR > 0.0001) elevated *CHRD, WNT5B* and *SHH*, consistent with a node-like expression profile, and relatively lower *SOX17* and *KRT18* than FNS+ cells (Fig 5e, Supplementary Table 6). We next looked for “mesendodermal” signatures within our groups, and only a small number of node-fated cells fit these criteria (FTS+) (Extended Data Fig. 8). We also observed divergent trends of gene co-expression within the cell groups (Extended Data Fig. 8c). *NODAL* and *EOMES* expression were negatively correlated with *FOXA2* in cells where *TBXT* was expressed (R <-0.15, p > 0.0001); the trend was the opposite in those that did not express TBXT (R >0.27, p > 0.0001). Accordingly, *NODAL* and *EOMES* were negatively correlated with *TBXT* (R <-0.35, p > 0.0001). Together these data further suggest that FT+/FTS+ cells contribute to the node/notochord. Given the APS origin of the node, and that all the cells in the APS/node trajectory do not express markers of classical EMT (Fig. 4d) this also suggests that the FTS+ cells are not mesendoderm. Instead, it appears that most cells that contribute to DE are *TBXT*- and correspondingly most mesoderm-fated cells are *FOXA2*-. We also find these observations apply in a less binary fashion to mouse embryos, with the high *T* population being node/notochord fated and a low *T* population, endoderm fated (Extended Data Fig. 9; Supplementary Table 7). Together this data demonstrates that the precursors of mesoderm, endoderm and node cells are temporally, spatially and transcriptionally distinct in the pig embryo.

### Extraembryonic signalling shifts correlate with the emergence of DE

Previous studies demonstrated the contribution of extra-embryonic endoderm to DE in mice^2, 39–41^. To establish whether this is true in pig embryos we isolated DE, gut, hypoblast and extra embryonic endoderm (ExE) clusters for further analysis (Extended Data Fig. 10a-c). Through marker expression and module scoring (See Methods) we identified subclusters of the hypoblast including the posterior hypoblast and AVE as well as distinct yolk-sac endoderm (Extended Data Fig. 10d-e; Supplementary Table 8). Temporal dynamics and module scoring also revealed a population of cells that appear to be an intermediate (InterHypo) between hypoblast and DE in line with similar observations in mice^39^ (Extended Data Fig. 10f-g). Isolation of E14-E15 samples further allowed for the identification of gut sub-populations that resemble their mouse counterparts^40^ (Extended Data Fig. 11; Supplementary Table 9). We also confirmed previous observations that suggested the hypoblast is the primary source of NODAL in pig embryos^42^ (Extended Data Fig. 10h-i). However, NODAL signalling in the hypoblast is restricted to E11.5-E12, which coincides with the specification of the DE from epiblast cells. Moreover, the absence of *NODAL* at E12.5 also correlates with the fate switching of the APS cells from endoderm to node/notochord. These findings corroborate recent reports suggesting that timed Activin/NODAL inhibition promotes notochord formation from DE-competent cells^43, 44^.

### Organiser-like signalling patterns of porcine cell types

It has been suggested that in mice, the node, PS and hypoblast have functions analogous to the organizer in amphibians^45^, yet it is unclear whether the signals leading to A-P patterning are relevant to other mammalian species. One such example is *Wnt3*, secreted by the posterior epiblast in response to *Bmp4*, which acts as the primary driver of mouse gastrulation^46^. Experiments in hESC and mESC have demonstrated that of the two *Wnt3* orthologues, *WNT3* and *WNT3A,* only WNT3 responds to BMP4^47^. Notably, humans with a homozygous WNT3 nonsense mutation display tetra-amelia and urogenital defects^47, 48^, a phenotype with a great resemblance to Wnt3a^-/-^ mice^49^. Further, the only demonstration of WNTs organiser function in large mammals comes from *in vitro* experiments using WNT3A^47^. Given the lack of understanding of the *in vivo* role of WNTs in large mammals we investigated canonical WNT crosstalk in E11.5-E12 cell types using CellChat^50^ (Extended Data Fig. 12a-b). In line with findings in mice and consistent with a role in A-P patterning, many early cell types were highly receptive to WNT ligands via multiple FZD and LRP receptor combinations. The highest *WNT3* signals came from the PS and APS (Extended Data Fig. 12a-b). Conversely, *WNT3A* was expressed within anterior ED clusters, namely the epiblast and APS. This previously undescribed WNT signalling pattern may explain why *WNT3A*-treated cells express GSC and can induce a secondary axis upon transplantation despite only WNT3 being downstream of BMP4^47^.

Cell-Cell signalling analysis also showed that multiple cell types were predicted to be receptive to node-produced SHH (Extended Data Fig. 12c). Notably, the scarcity of node cells identified prior to E12.5, coupled with the fact that *NODAL, CER1, LEFTY2* and *DKK1* were produced by DE but not node-fated APS cells (Fig. 6e, Supplementary Table 6), suggests that the mammalian node/notochord is principally involved in secondary gastrulation^45^. Consistent with this hypothesis is that node cells and node-fated anterior streak cells not only secreted SHH, a dorsal tissue patterning ligand, but also expressed high levels of *FGF8, NOG* and *CHRD*, factors primarily associated with axial extension and DV patterning^51–53^.

### A balance of WNT and NODAL signalling determines the differentiation of DE from epiblast cells

WNT and NODAL signalling are critical for endoderm formation^47^ however, the balance of these competing signals required for DE formation has not been fully established. To assess this, we used varying concentrations of Activin A (ActA) and a WNT agonist (CHIR99021, CHIR) to simulate specific microenvironments across the ED in 2D cultures of the pig EDSCs line (EDSCL4)^21^ and hESC lines H9 and HNES1^54^. All cells were cultured in AFX medium (see methods). A dose-response effect was observed with the number of FOXA2 positive cells increasing with higher ActA levels at both 24 and 48 hrs in EDSCs, though this was partially stunted by increasing levels of CHIR. In both human and pig stem cells SOX17 increased by 48 hrs when exposed to high levels of ActA (100ng/ml) in the presence of endogenous WNT or CHIR. In contrast, when endogenous WNT was inhibited using XAV939, SOX17 was very low, indicating a requirement for low levels of WNT signalling (Fig 6. a-b, Supplementary Data Fig. 1). A similar response was determined for human cells, but a reduced threshold of ActA was needed for peak FOXA2 and SOX17 expression. In line with recent reports, TBXT expression required exogenous Activin/NODAL, in addition to WNT, suggesting a role for the hypoblast in streak formation. In both species, higher levels of CHIR increased TBXT expression at 24 hrs, which was largely extinguished by 48hr, consistent with the expression of TBXT in the embryo. A small number of SOX17+ only cells could be observed from 8 hrs after treatment. Co-expression between SOX17 and TBXT or SNAI1 was rarely observed even at 24-48 hrs and showed very low Pearson’s correlation scores (Supplementary Data Fig. 2a-e).

Based on these results we propose a model of DE formation in the pig embryo (Fig. 7c) whereby between E10.5 and E11, the first TBXT+ cells emerge at the posterior ED opposing the anterior hypoblast/AVE-produced WNT/NODAL inhibitors (*LEFTY* and *DKK1*)(Extended Data Fig. 11d), consistent with previous work in pig^42^. Driven by a combination of posterior epiblast-secreted WNT and hypoblast-derived NODAL signalling, these cells initiate EMT, thus constituting the nascent mesoderm. The interaction of reduced WNT signalling and increased *NODAL* induces *EOMES* in cells anterior to the TBXT ED domain. *EOMES*, which is capable of inhibiting *TBXT* activity, alongside *NODAL*, promotes *FOXA2* expression. The most anterior FOXA2+ cells produce the NODAL inhibitor *CER1*, delaminate and intercalate into the hypoblast to form and expand the DE. This results in the termination of NODAL, but not WNT signalling in the newly formed DE. The remaining FT+ NODAL-primed cells, still driven by WNT signalling, begin to form the node from approximately E12 onward.

**Fig. 7.**
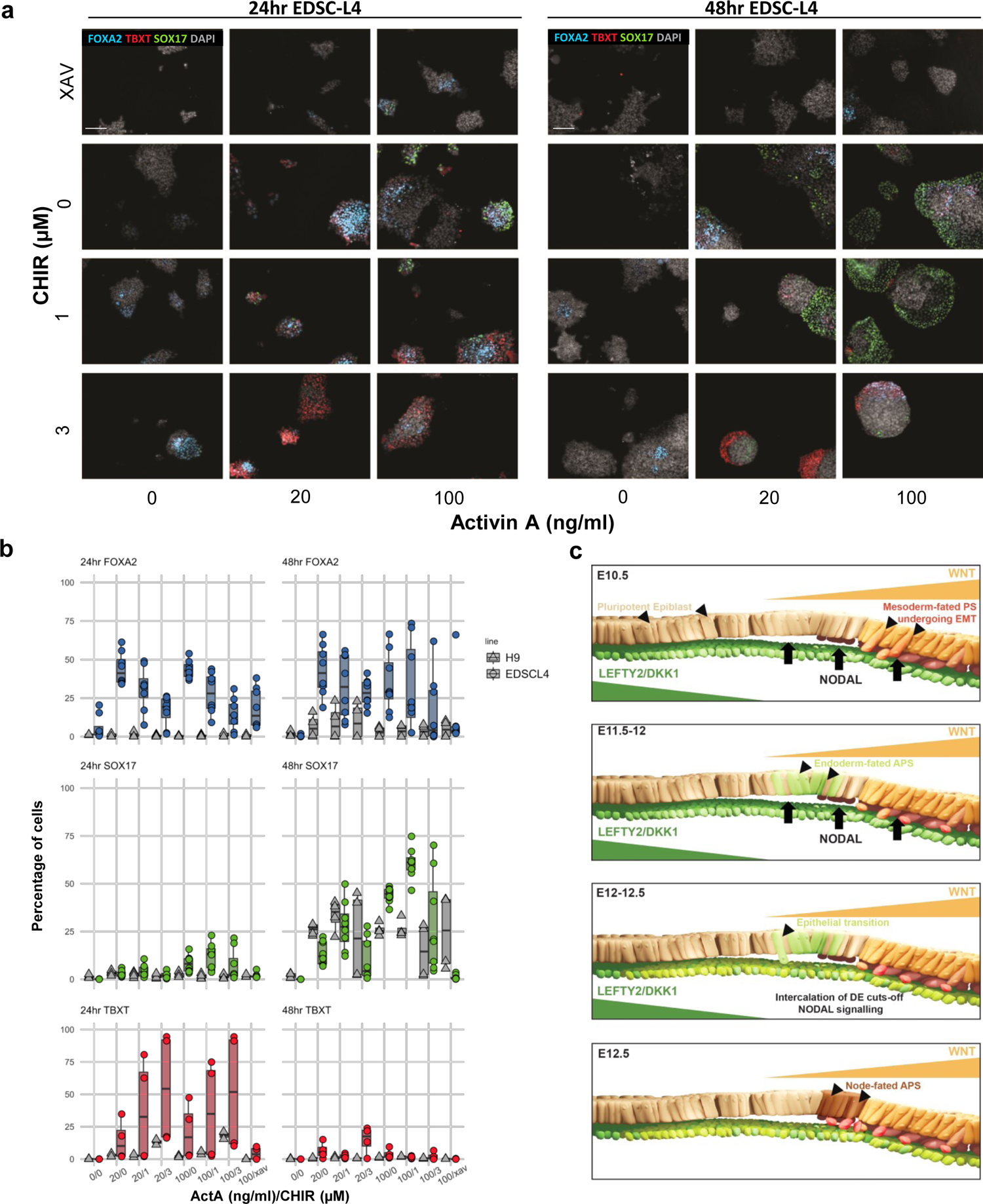
Effect of ActA and WNT in pig EpiSC and hESC. **a,** Representative images depicting differentiation conditions for EDSCL4. Images were captured using an Operetta CLS high-throughput microplate imager. Scale bar: 200 µm. **b,** Box plot summarizing 2D differentiation experiments. Data is normalised to well background signal and results represents biological triplicates. **c,** Proposed model of epiblast-DE differentiation in pig embryos (see text for details). anterior primitive streak (APS).

## Discussion

We present a scRNAseq atlas of pig gastrulation and early organogenesis that represents a comprehensive resource for exploring the molecular mechanisms governing cell-fate determination during a crucial juncture of development. We harnessed this resource to investigate the major temporal signalling and differentiation events which direct primary gastrulation in bilaminar disc embryos. Our findings underscore the nuanced heterochrony in embryonic development across mammals, evidenced by asynchronicities in cell type maturation. Further, these heterochronic differences combined with comparisons between pigs, monkeys and mice further reinforce the idea that extraembryonic tissues may be less conserved than their embryonic counterparts^3, 27, 29^. Through cross-species mapping and comparative transcriptomics, we also elucidated distinctive patterns of gene expression associated with key signalling and adhesion pathways in pig and monkey epiblasts, compared to their mouse counterparts. Such observations may have implications for understanding species-specific aspects of cell differentiation, growth, and morphological features.

Despite the observed species differences, we were for the first time able to identify highly conserved early cell-type specific programs between mice, primates and pigs. These findings were exemplified by our investigations into the segregation of the endoderm, mesoderm and node, the precursors of which display organizer-like patterns of gene expression. With the reduced spatial constraints and slower development of the pig embryo compared to mice, we showed a distinct temporal and spatial pattern of DE and specification from ED cells via an epithelial-to-epithelial transition a model which had previously only been demonstrated conclusively in mouse models^15, 16, 31^. Further, our findings extend the understanding of the emergence of the node in mammals, which contrary to chick models^45^ and earlier DE progenitors, we found no evidence for a role in A-P patterning. Combined with both embryo whole mount IF and our *in vitro* experiments we showed that this sequence is determined by a balance of WNT and Activin/NODAL gradients along the A-P and D-V axis, respectively, and equivalent findings were recapitulated *in vitro* with hESC, suggesting that these findings are representative of human embryos.

We anticipate that this new resource in combination with other recent works^3^ and novel techniques, such as spatial transcriptomics of early embryos^55^, will inform more detailed analyses of species differences and will shed light on the molecular events that underly the phenotypic differences in mammalian embryos. Given their use in agriculture pig embryos represent a highly accessible model for functional investigations relevant to human development. As with comparable datasets in mice^2^, we expect that this dataset will serve as a wild-type reference for comparisons against mutant embryos. Such investigations will serve as a foundation on which to develop more robust in vitro differentiation protocols of pluripotent cells, novel approaches to interspecies chimaeras and to study the development of organ used for xenotransplantation^8–10^.

## Methods

### Ethical statement

All procedures involving animals were approved by the Animal Welfare and Ethics Review Committee (Nbr. 99) of the School of Biosciences, The University of Nottingham. The research conducted adhered to the Home Office Code of Practice guidelines for the Housing and Care of Animals used in Scientific Procedures.

### Embryo collection

Embryos were collected from crossbred Large White and Landrace sows (2–3 years old) between days 10 to 16 post artificial insemination. Each uterine horn was separated into an upper and lower half before fresh PBS+5% BSA was used to flush out porcine embryos. Uterine horns were then bisected and searched by hand for any further embryos. Embryos were stored in warm PBS+5% BSA during. Embryos were either fixed in PFA at 4°C overnight for IHC or taken for 10x single cell RNA sequencing.

### Immunohistochemistry and imaging of whole mount embryos

Embryos were permeabilized and blocked at the same time for 2hr in a solution containing 10% donkey serum and 5% BSA with 1% Triton-X (PB buffer) at RT. Samples were incubated with primary antibody O/N at 4°C in PB buffer. Secondary antibodies were incubated with sample for 2hr at RT in PB buffer. Washes were performed after primary and secondary antibody incubations 4×15min in PBS with 0.2% Triton-X. Samples were mounted in either VECTASHIELD or Fluoroshield with or without DAPI. If the mounting media did not contain DAPI, it was added at the secondary antibody incubation stage. Antibodies used here are listed in Supplementary Table 10. Imaging of embryo samples was performed with a confocal Zeiss LSM 900 with Airyscan, all samples were imaged as z-stacks with a Z resolution of 0.32 µm.

### Segmentation, quantification and in silico recapitulation of embryos

Z stack images were segmented using a StarDist/TrackMate pipeline within Fiji. StarDist allows for 2D segmentation, and TrackMate is used to build up each cell in 3D from the totality of the 2D segmentation data. 3D data for each cell was exported as a .CSV file and further analysed in R using custom code. Analysis in R allowed for extraction of location and protein expression information for each cell, which was then used to recapitulate the embryos *in silico*, as well as directly quantify the number of cells expressing a given protein. Lateral re-slicing performed within Fiji.

### Culturing, imaging and quantification of 2D culture experiments

Both human and pig cells were cultured in AFX medium: N2B27 supplemented with FGF2 (20 ng/ml) + Activin A (12.5 ng/ml) + XAV939 (2 μM)^21^. For experimental procedures, cells were seeded at a range of seeding densities (750-1200 cells/well) into CytoOne TC treated 96 well plates coated with Laminin-511-E8 fragment with additional ROCKi for 48hrs. Cells were differentiated in N2B27 + CHIR, Activin A and XAV. Cells were then fixed with 4% PFA and immunostained as detailed above. Images were captured using an Operetta CLS high-throughput microplate imager and segmented using Harmony (v4.1) from phenoLOGIC. Thresholding, cell expression identity and plot making was performed with a custom script in R. Cell lines used were EDSCL4 (porcine), H9 and HNES1 (human) (Table 10).

### Single-cell transcriptomic analysis

#### Preparation of scRNA-seq library and sequencing

Single-cell libraries were constructed using Single Cell 3 Library & Gel Bead Kit v3 according to the manufacturer’s protocol (10X Genomics). Briefly embryos taken for scRNAseq were either frozen and stored at −20C or sequenced fresh. Given the large size of ExE tissues, to ensure that embryonic cells were well represented we manually dissected most ExE structures prior to dissociation and sequencing, preserving the hypoblast. Embryos were dissociated into single cells via incubation with TrypLE, for 7 mins at 37°C. Embryos were further dissociated via pipetting and then washed with a solution of DMEM/F12 0.04%BSA to quench TrypLE activity. Cells were strained though a Flowmi cell strainer into a 15ml falcon. Once dissociated, cells from pools of same stage embryo were counted using a haemocytometer. Approximately 4000-14000 cells per lane of each 10x chip were transferred. The chip was then loaded into a Chromium Controller for cell lysis, cDNA synthesis and barcode labelling. cDNA libraries were assessed using an Agilent 2100 Bioanalyzer (Agilent Technologies). Finally, the libraries underwent 150bp paired-end sequencing using the NovaSeq 6000 platform.

#### scRNA-seq data preprocessing

Raw fastq files were processed using cellranger-6.1.2 software with default mapping arguments. Reads were mapped to the Sus scrofa Sscrofa11.1 (GCA_000003025.6) genome. Subsequently, the CellRanger ‘aggr’ command was used to normalize the sequencing depth of different samples. For samples where the number of sequenced cells differed greatly from the counted number of cells prior to loading the ‘force cells’ command was used, and the estimated number of loaded cells was given.

#### Filtering of cells, integration, dimensionality reduction and clustering

The filtered expression matrix with cell barcodes and gene names was loaded with the ‘Read10X’ function of the Seurat (v.4.0.0) R package^56^. Following this, Seurat objects from each sample (23 samples) were independently created and processed according to standard Seurat protocols. Initially, single cells with a number of detected genes (nFeature_RNA) above 1750 were retained to exclude low-quality cells. Subsequently, doublet or multiplet cells were identified with the DoubletFinder R package^57^ and excluded. After normalization of the Seurat object, we selected the 3000 most variably expressed genes using the ‘FindVariableFeatures’ command. These features were used to calculate the first 100 principal components. Given the heterogeneity of cell compositions of early and late embryos, we did not exclude cells based on the percentage of their transcriptomes that were made up of mitochondrial genes. Instead, following clustering we looked for non-discreet clusters that had significantly higher expression of mitochondrial genes and low ‘nFeature_RNA’. Two clusters of cells were excluded on this basis, we also noted that these cells showed expression of markers of multiple cell-types suggesting that these cells were clustering based on a shared apoptotic identity. We then used the ‘FindIntegrationAnchors’ and ‘IntegrateData’ functions of Seurat to exclude individual heterogeneities between samples. Data integration was done using the reciprocal pca (rpca) method and the first 30 dimensions of each object were used. To construct the main UMAP plot of 91,232 cells in Fig 1c, we used the first 25 principal components for calculating UMAP 1 and 2, setting the seed at 42, using a minimum distance of 0.5 and the 50 nearest neighbours (n.neighbors). The other parameters were kept as the defaults for UMAP generation. For clustering of the same cells, the ‘k.parameter’ of 20 and ‘n.trees parameter’ of 50 were the default settings during the neighbour-finding process; 25 dimensions were selected via the ElbowPlot method for neighbour finding, and clustering. A resolution of 1.2 was used to identify the 36 major cell-types. For sub-clustering, cell-types of interest were subsetted, objects were re-scaled and then the same numbers of principle components were calculated. The same parameters were used for neighbour finding, however, a resolution of 0.5 was used to clustering. For UMAP creation, again the ElbowPlot method was used to select the number of dimensions however the 30 nearest neighbours were used along with a minimum distance of 0.4. We calculated the DEGs of each cell cluster with RNA assay using the ‘FindAllMarkers’ function in Seurat. Heatmaps were plotted based on the most highly expressed genes (according to fold change) which had an adjusted p-value less than 0.05.

#### Pseudotime

The ‘monocle2’ R package^58^ was used to calculate the developmental pseudotime of cells. The Seurat object was converted to a monocle2 object by the ‘as.CellDataSet’ command of the SeuratWrappers R package. The ordering filter was then constructed using the results of ‘FindAllMarkers’ using a ‘min.pct’ of 0.5, a ‘logfc.threshold’ of 0.5, a ‘return.thresh’ of 0.01, and negative markers were excluded. The ‘reduceDimension’ command was used with the ‘DDRTree reduction method. Lastly, the ‘orderCells’ command was used to analyse developmental pseudotimes. Pseudotime trajectories were visualized using the ‘plot_cells’ function.

#### Gene expression-based categorisation of cells

Cells were categorised as having ‘positive’ expression of *NANOG, FOXA2, TBXT* or *SOX17* cells if they had a scaled expression value greater than 0.1.

### Transcription factor regulon analysis (SCENIC)

Gene regulatory networks and regulons were elucidated using the command-line interface (CLI) of the pySCENIC pipeline, including tools such as ‘arboreto_with_multiprocessing.py’, ‘ctx’, and ‘aucell’^59^. Input data consisted of raw gene counts, pre-processed to consider cells detecting between 1,750 to 7,500 genes. Cells bearing over 10% mitochondrial reads, along with genes identified in less than three cells, were excluded from the analysis. For transcription factor binding motif analysis, a custom RcisTarget database was assembled using the ‘create_cistarget_motif_databases.py’ tool. The construction of this database was based on the v109 Ensembl release of the Sscrofa11.1 reference genome. Feather ranking databases were constructed based on two distinct sets of regions: (1) regions encompassing 2,500 bp upstream and 500 bp downstream of each transcription start site (TSS), and (2) regions spanning 10 kb upstream and 10 kb downstream of each TSS. The binding motif list was generated by renaming human binding motifs, obtained from the SCENIC motifs’ v10 public collection (https://resources.aertslab.org/cistarget/motif_collections/v10nr_clust_public/snapshots/), to their pig orthologues. High-confidence, one-to-one orthologues were obtained using the Biomart tool available on the Ensembl website.

### Cell–cell communication analysis

Cell annotation information and raw count expression matrix was exported from Seurat, and pig gene names were converted to their human orthologues using the same process as described above. The matrix was then processed according to the standard CellChat^50^ protocol using the CellChatDB.human ‘secreted signalling’ database.

### Comparison of single-cell transcriptomic datasets among mice, humans, monkeys and pigs

To project pig single-cell data onto the mouse, human and, monkey datasets, expression matrices were exported from Seurat and gene names were converted to their human orthologues. Ensembl biomart was used to identify 14,108 high-confidence one-to-one orthologues. All genes that were not present in all four species were excluded from the matrices. Individual matrices were then loaded into Seurat and processed individually as before. Pig, mouse and monkey datasets were then randomly down sampled so that each sample contained 25,000 cells. The transferal of cell labels between each dataset was done in a pairwise fashion using the MapQuery function in Seurat. The anchors between mouse and pig data were found with the FindTransferAnchors function (reference.reduction, ‘pca’; dims, 1:50; k.filter, NA), and the function MapQuery (reference.reduction, ‘pca’; reduction.model, ‘umap’) was used. For comparisons of cell transcriptomes datasets were integrated using the IntegrateData function as before, cells were then rescaled and the active identity was set to the pig cell type annotation, three individual objects were made from this parent object containing pig-mouse, pig-monkey and mouse-monkey. The ‘FindConservedMarkers’ function was used to find conserved and divergent cell type-specific markers. A marker gene was classified as ‘conserved’ if the marker was significantly increased beyond 2-fold in the cell type of interest compared to all other cell types (adjusted p-value <0.05).

## Data availability

The dataset of pig gastrulation and early organogenesis generated in the current study is available in the NCBI Gene Expression Omnibus (GEO) under accession no. GSE236766. The mouse gastrulation and early organogenesis dataset used as a reference is available at ArrayExpress under accession no. E-MTAB-6967. The monkey gastrulation dataset is available at GEO, under accession no. GSE193007. The human CS7 dataset is available at ArrayExpress under accession no. E-MTAB-9388 and at GEO under accession no. GSE157329. Source data are provided with this paper.

## Acknowledgements

This project was supported by the Biotechnology and Biological Sciences Research Council grants [grant number BB/S000178/1] [grant number BB/T013575/1] to R.A, M.L, and J.N. We thank the staff at the nMRC core imaging at the University of Nottingham, especially Jacqueline Hicks.

## Author Contributions

L.S, A.S, B. P and S.K designed and performed experiments including scRNA-Seq, embryo dissections and wrote the paper; L.S, A.S, S.H, F.S, D.G and T.L performed bioinformatic analysis; A.S., T.A and D.K. performed IF and functional experiments; M.L. supervised scRNA-seq analysis; J.N. supervised the functional experiments and wrote the paper. R.A. supervised the project, designed experiments, performed dissections, and wrote the paper. All authors discussed the results and contributed to the manuscript.

## Competing Interest

The authors declare no competing interests.

---

**Extended Data Fig. 1.**
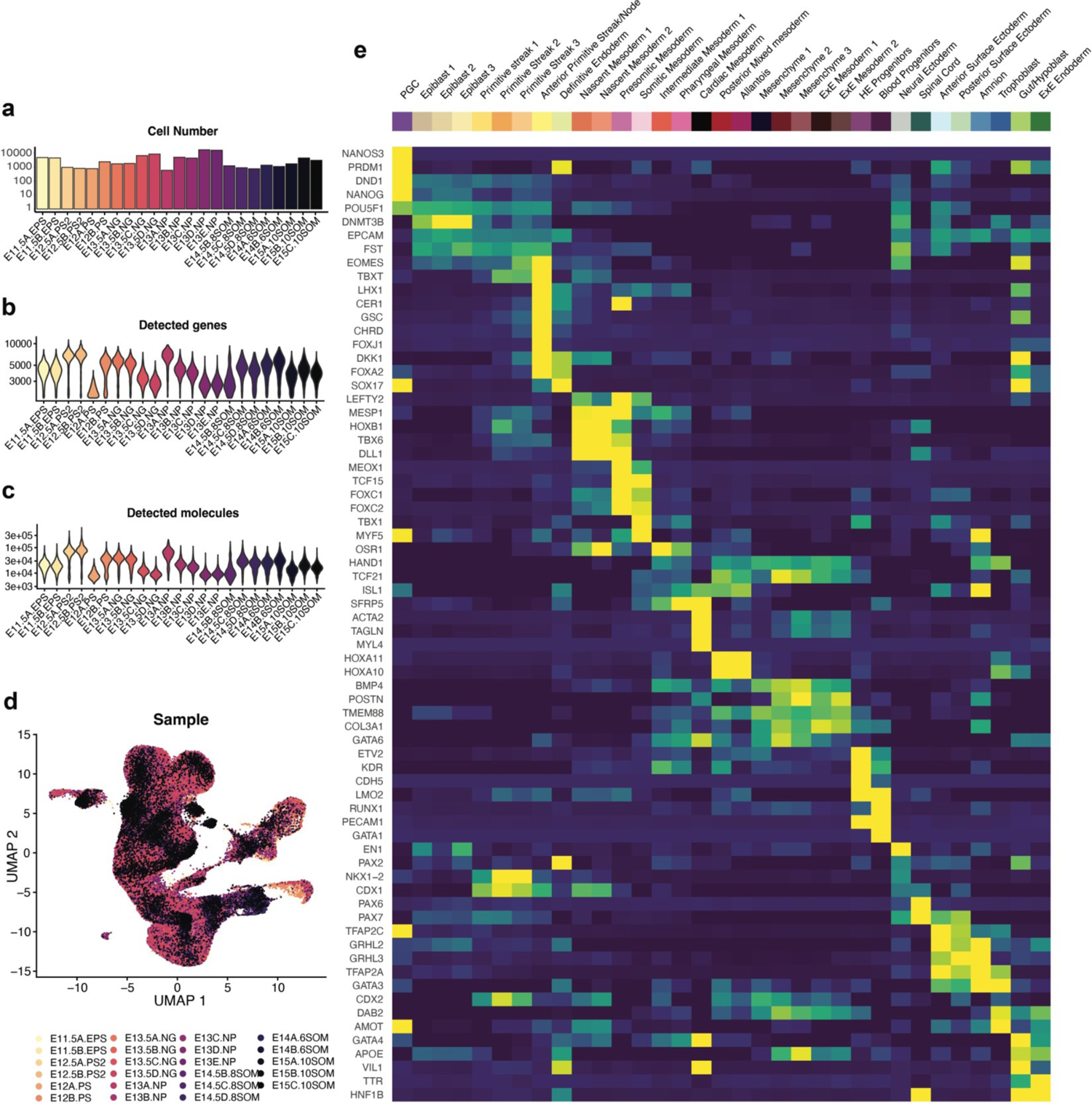
Cell distribution per sample and gene markers used to define cell identity. **a**, Number of cells captured in this study in each sample used in this atlas. **b**, Violin plots illustrating the number of detected genes per cell per sample. **c**, The number of molecules detected per cell per sample. **d**, UMAP plot showing atlas cells from Fig.1c) coloured by sample, showing consistency across replicates from the same stage. **e**, Heatmap showing the markers used for cell type identification, a full list of these markers and the source publications are available in Supplementary Table 1. ExE; Extra-embryonic, HE, Haematoendothelial.

**Extended Data Fig. 2.**
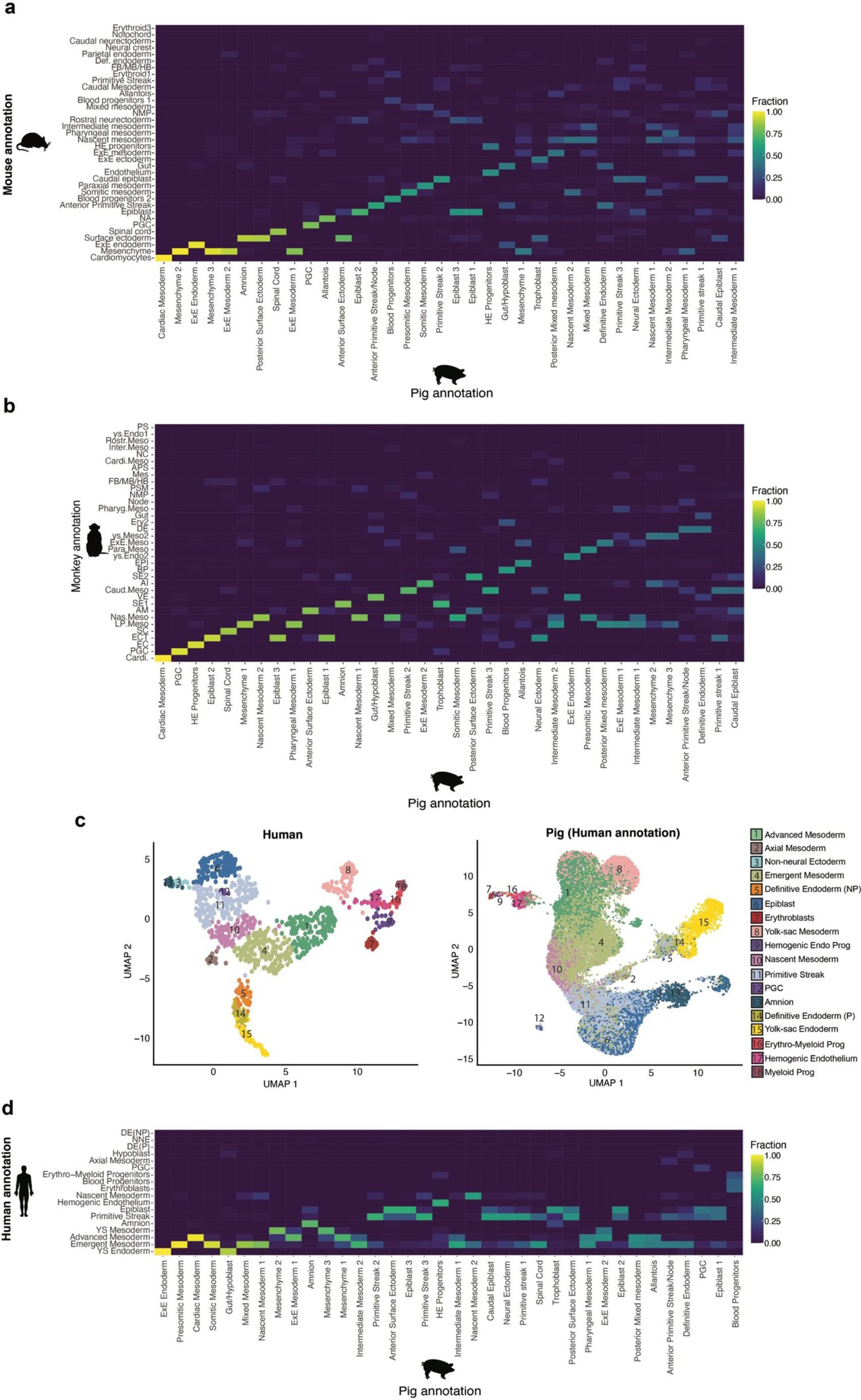
Comparisons of mouse, monkey and human cell annotations. **a**, Heat map showing the fraction of pig cells grouped by cell type allocated to a particular mouse cell type after label transfer. **b,** Heat map showing the fraction of pig cells grouped by cell type allocated to a particular monkey cell type after label transfer. **c**, UMAPs showing E17-19 human embryo cell types (left)^30^ and pig E11.5 to E15 with mouse annotations (right) after reciprocal PCA-based projection onto the human dataset. **d**, Heat map showing the fraction of pig cells grouped by cell type allocated to a particular human cell type after label transfer. A fraction of 1 indicated that 100% of cells of a given cell type were predicted to be analogous to the cell identity in the queried organism.

**Extended Data Fig. 3.**
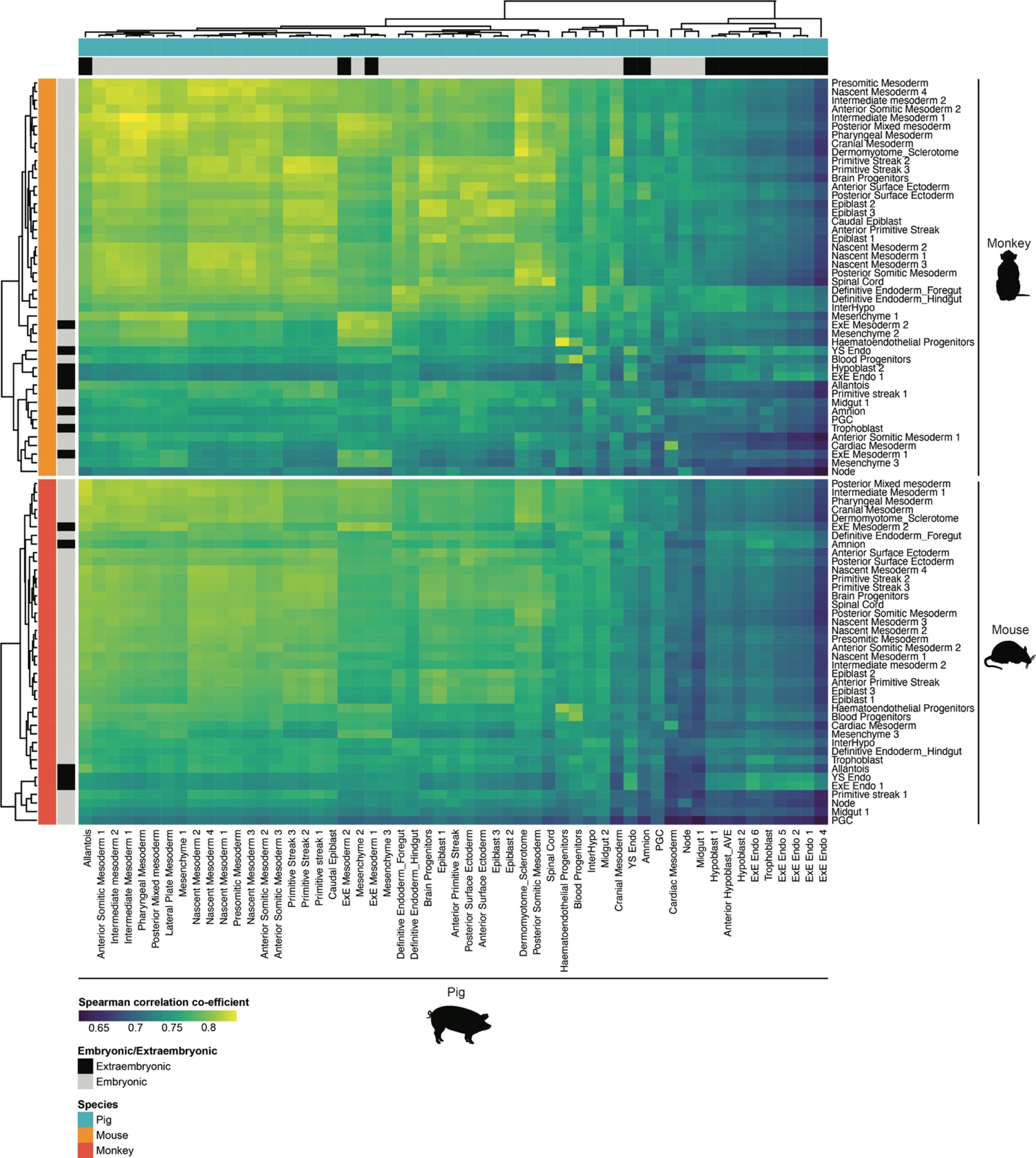
Extra-embryonic cell types show lower correlation across mammals. Heatmap showing Spearman correlation coefficient between the transcriptomes of matched cell types in pig, monkey and mouse embryos.

**Extended Data Fig. 4.**
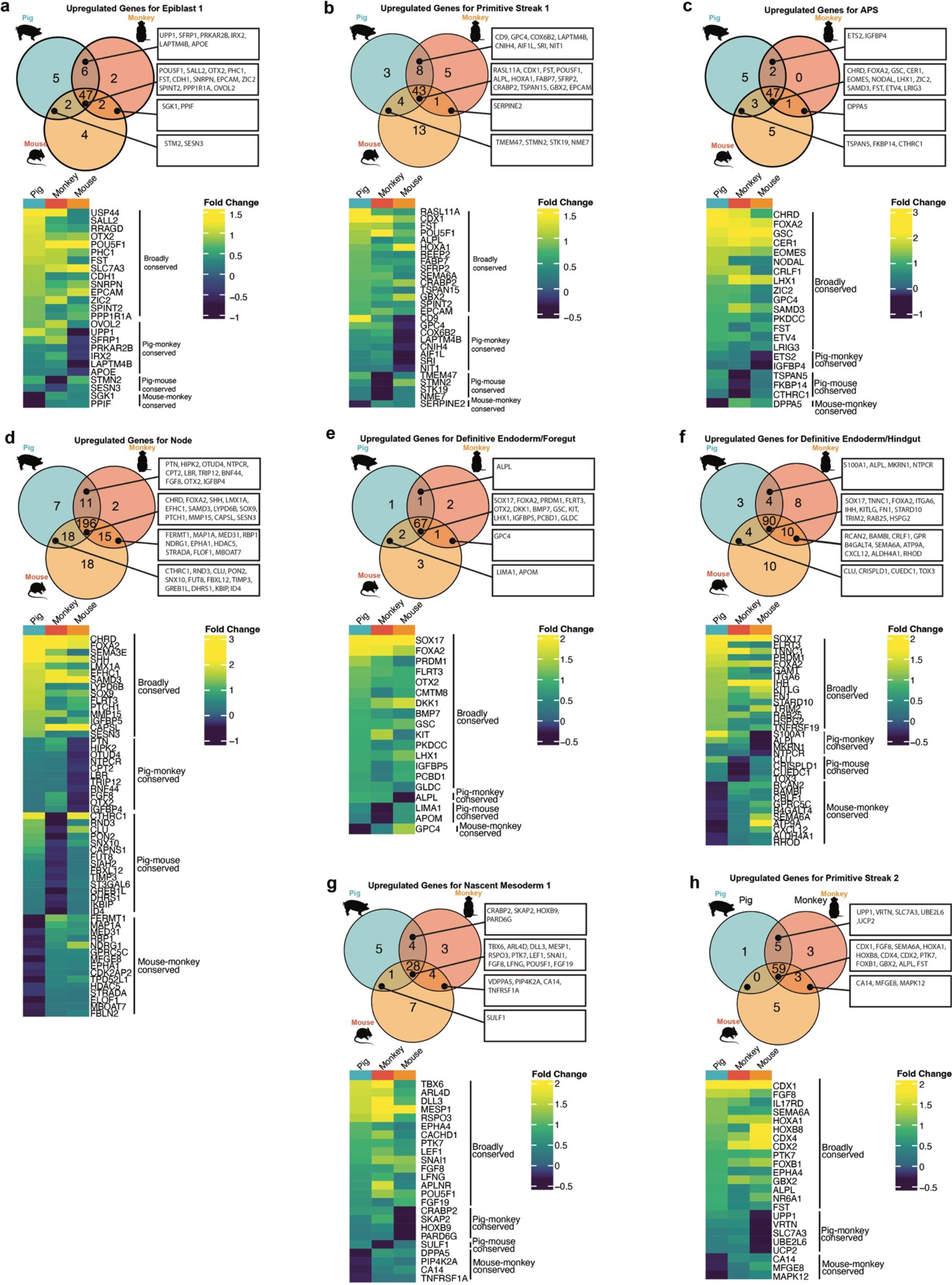
a, Cell type-specific gene expression programs are highly conserved. **a-h,** Venn diagrams and heatmaps showing unique and overlapping cell type-specific genes in pig, monkey and mouse. The heatmaps illustrate the average fold change in gene expression within the cell type of interest, compared to the mean fold change across the remaining cell types for a specific species. APS, Anterior primitive streak.

**Extended Data Fig. 5.**
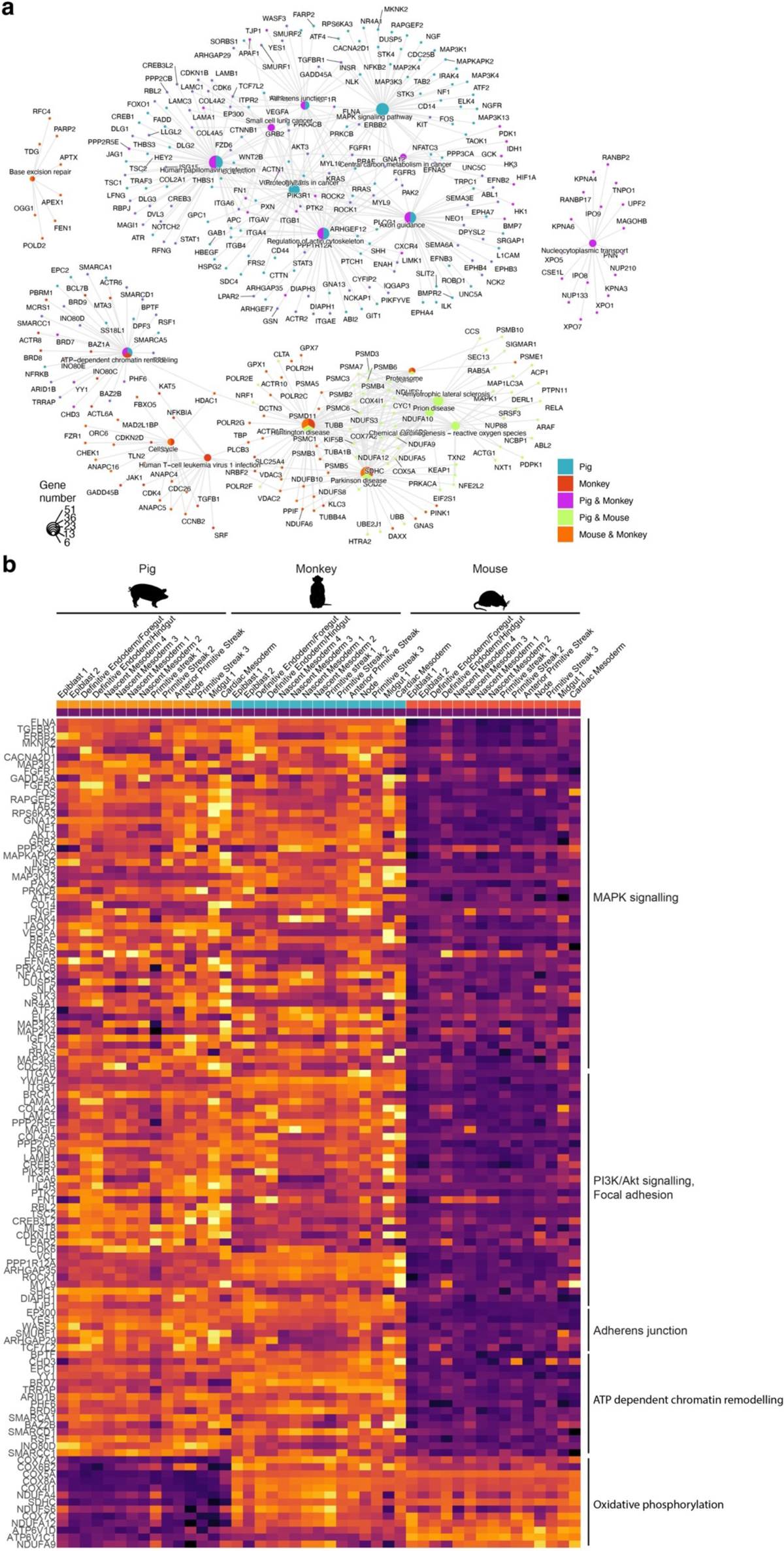
a, Broad differences in early cell behaviour are observable across mammals. **a,** Plot of Cnet illustrating the enriched KEGG terms and associated genes found in both unique and shared expression profiles among pig, monkey, and mouse epiblast cells. **b,** Heat map of selected KEGG terms-associated gene expression in early pig, monkey and mouse cell types.

**Extended Data Fig. 6.**
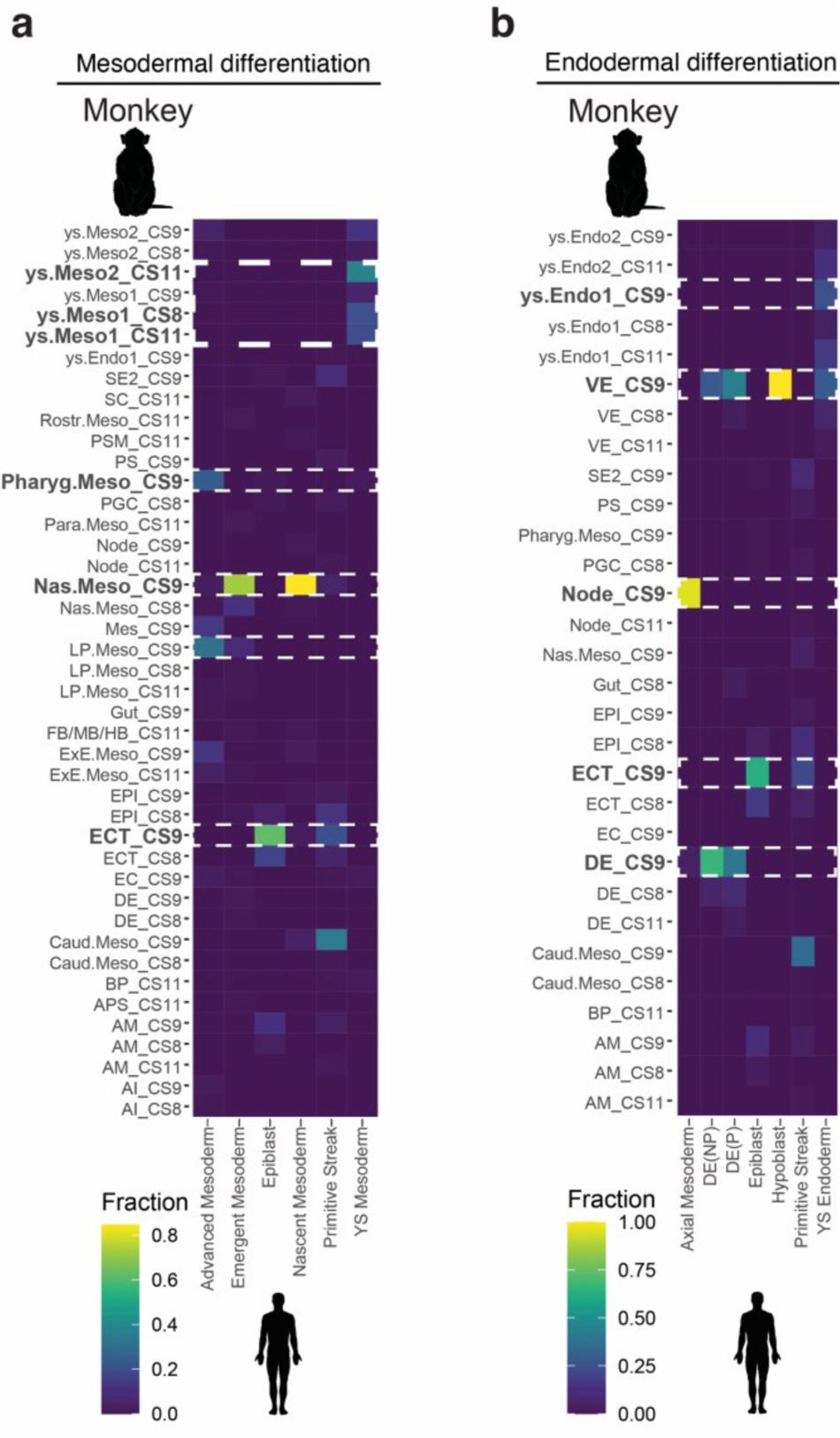
**a**, Homochronicity among monkey and human cell type differentiation Heat maps showing the fraction of human mesodermal cells^30^ grouped by cell type allocated to a monkey cell identity^6^ after label transfer. **b,** As with a, except with endodermal cell types. A fraction of 1 indicated that 100% of cells of a given cell type were predicted to be analogous to the cell identity in the queried organism.

**Extended Data Fig. 7.**
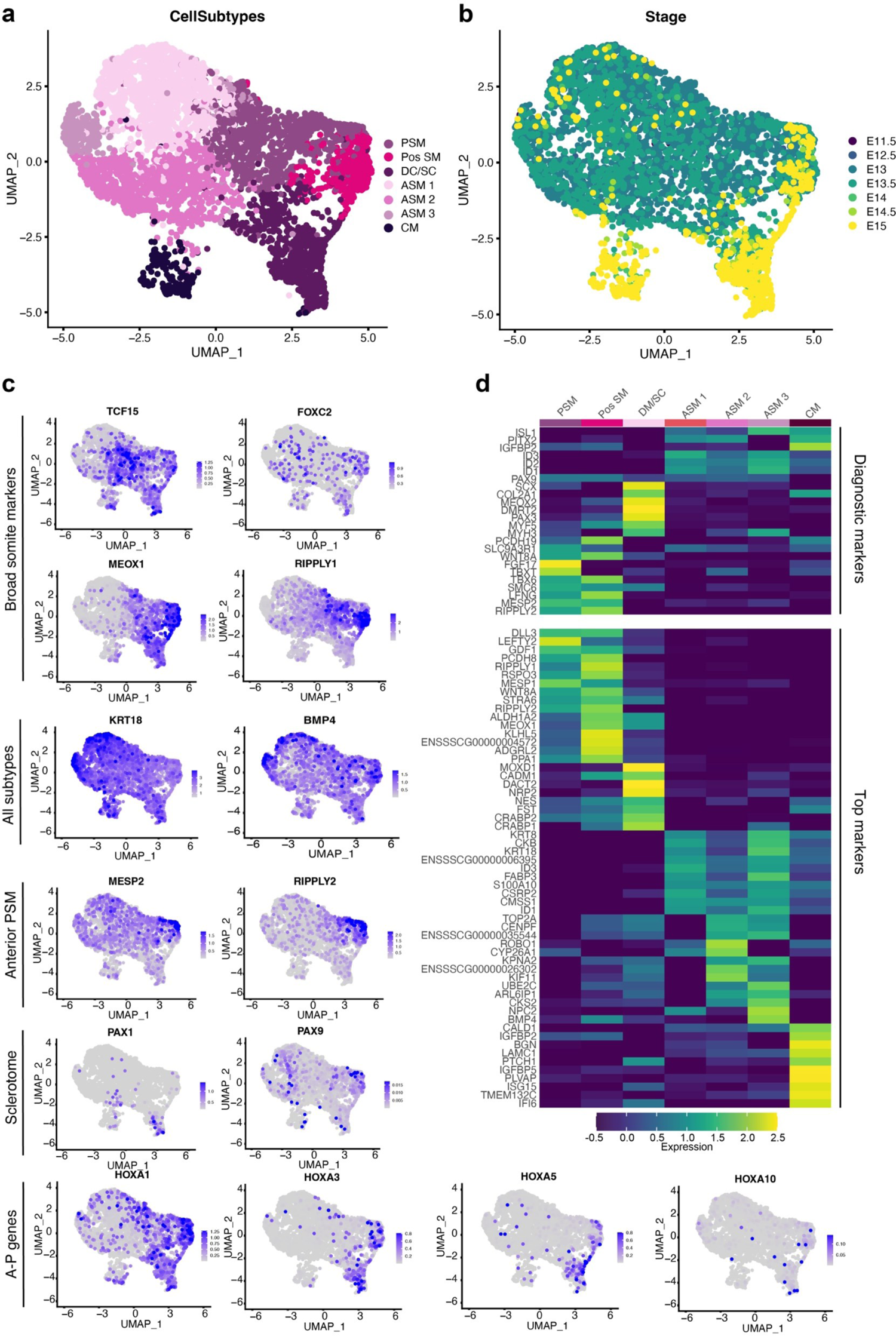
Somitogenesis in porcine embryos. **a**, UMAP plot showing sub-clustering of Presomitic and Somitic mesoderm clusters. 7 subclusters were identified: Presomitic mesoderm (PSM), Posterior somitic mesoderm (Pos SM), Dermomyotome/Sclerotome (DC/SC), Anterior Somitic Mesoderm 1, 2 and 3 (ASM1, ASM2, ASM3), as well as Cranial Mesoderm (CM). **b**, UMAP plot showing cells from a. Cells are coloured by embryonic day. **c**, UMAP feature plots of key markers of somite development. **d**, Heat map illustrating the scaled average expression of selected genes. Top: diagnostic marker genes for each subcluster. Bottom: Top 10 most significant marker genes.

**Extended Data Fig. 8.**
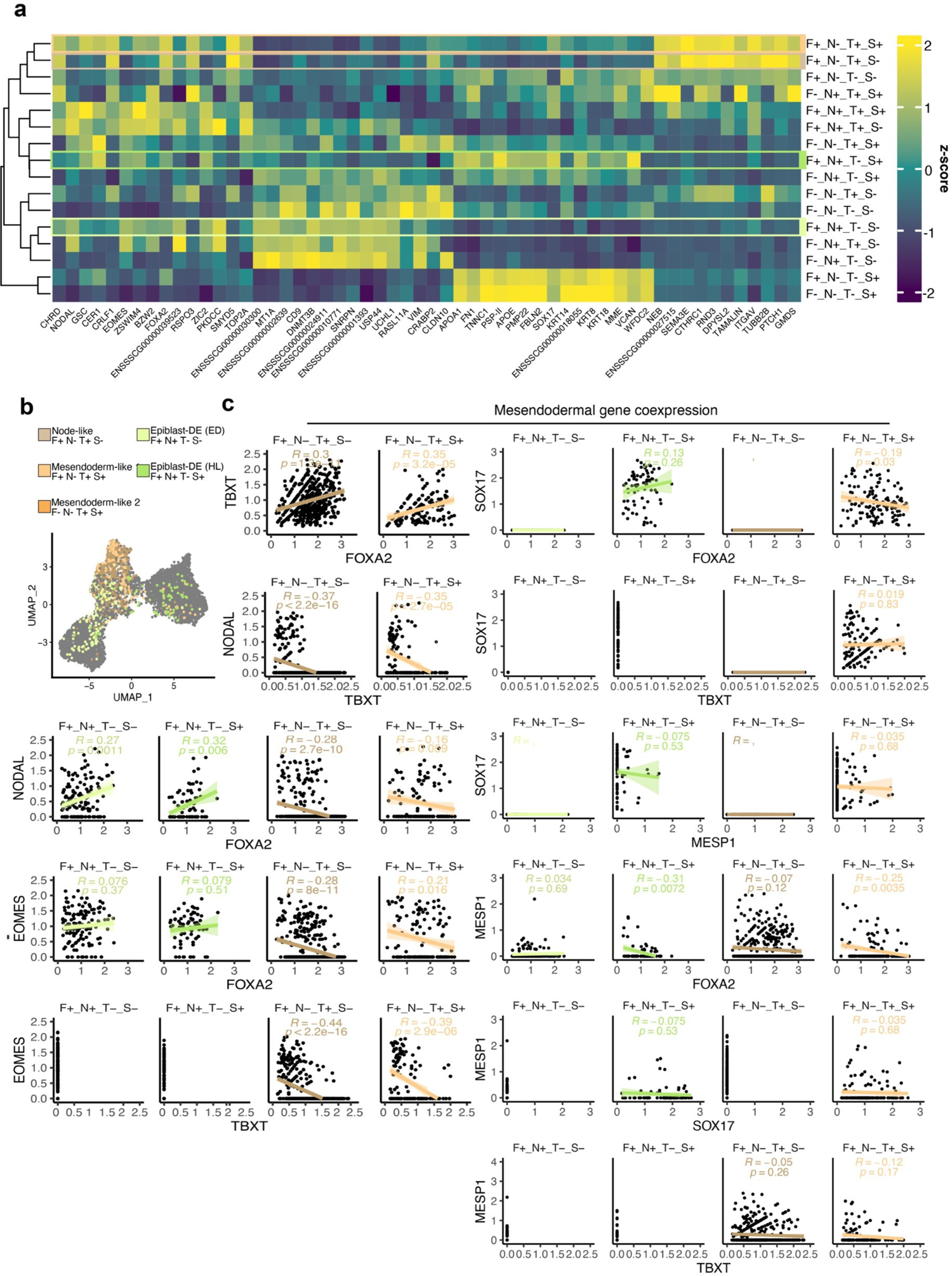
Mesodermal and endodermal genes are mutually exclusive. **a**, Heat map illustrating the scaled mean expression of the top marker genes for F/N/T/S categorised cells from Fig 5.a & Extended Data Fig.2a. Only the top 20 most significant genes for each subcluster are shown. Column clustering indicates similarity. Legend/key shared with b&c. **b**, UMAP plot categorised by FOXA2, NANOG, TBXT and SOX17 expression. F, FOXA2; N, NANOG; T, TBXT; S, SOX17 cells. Cells are coloured by their F/N/T/S category. **c**, Scatter plots showing mesoderm and endoderm associated transcription factor co-expression of F/N/T/S categorised cells from Fig 4.a & Extended Data Fig.2a. Correlation co-efficient (R) and adjusted p-value (p) following Pearson correlation test indicated.

**Extended Data Fig. 9.**
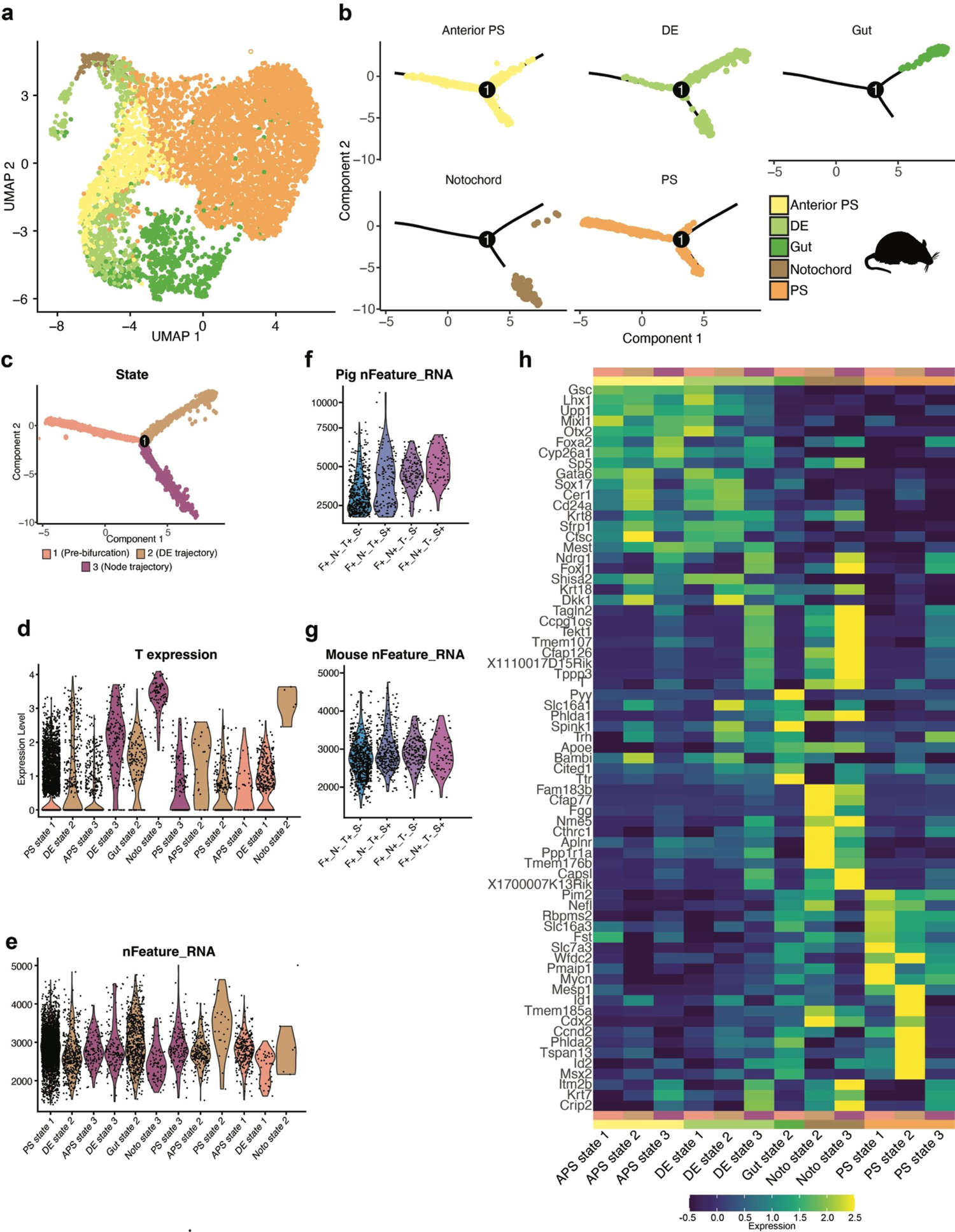
Endoderm forms from TBXT-low cells in mice. **a**, UMAP plot showing PS, APS, Notochord, DE and Gut clusters (5,860 cells) from Pijuan-Sala et al^2^, coloured by cell type. **b**, Force-directed graph of the cells illustrated in a, after pseudo-temporal ordering using Monocle2. Force-directed graphs are split by cell type. **c** Force-directed graph of the cells illustrated in a and b coloured by state. State 1 represents cells that have not committed to notochord or DE/Gut fates. State 2 represents cells moving toward a DE/gut fate. State 3 represents cells moving toward a Notochord fate. **d**, T expression in the cell types shown in a split into their respective differentiation states defined in c. **e**, Violin plots showing the number of mapped genes in each cell type defined in d. **f**, Violin plots showing the number of mapped genes in the cell types of interest defined in Figs 5-7. **g**, Violin plots showing the number of mapped genes in the mouse-equivalent cell types shown in f. Both mouse and pig DE/Node progenitors show a comparable number of mapped genes. **e**, Heat map illustrating the scaled mean expression of the top 10 most significant DEGs between the cell types from c-e.

**Extended Data Fig. 10.**
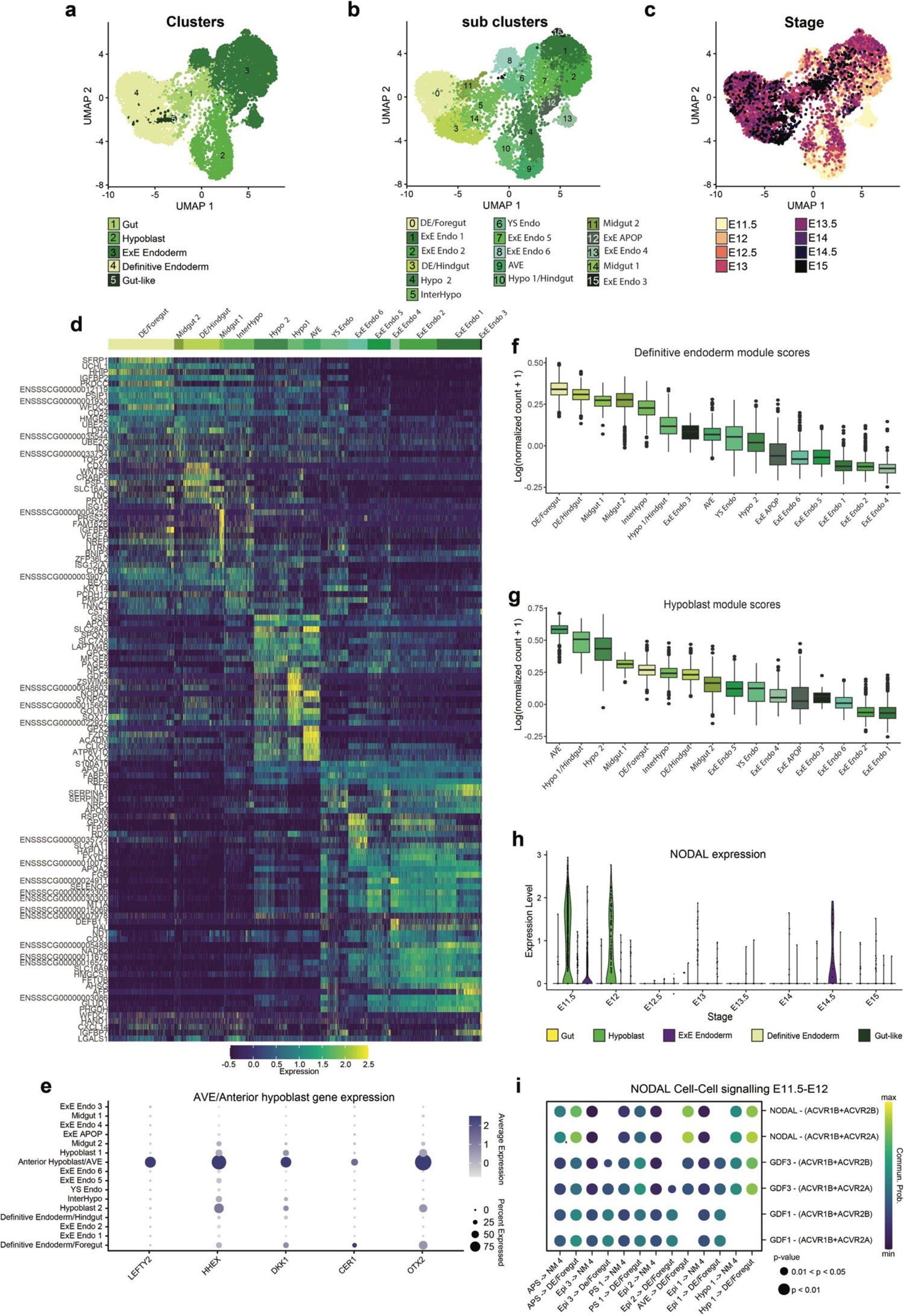
The hypoblast is a major source of signalling and contributes to DE. **a**, UMAP plot showing DE, gut-like, gut, hypoblast and ExE Endoderm subclusters coloured by cell-type. **b**, UMAP plot showing embryonic and extraembryonic sub-clusters coloured by cluster. **c**, UMAP plot showing cells from a&b coloured by timepoint, showing stage specificity of clusters. **d**, Heat map illustrating the cluster-scaled mean expression of the top 10 most significant newly discovered marker genes for each subcluster. **e,** Dot plot showing the average expression and percentage of cells expressing AVE/Anterior hypoblast related genes. **f&g**, Box and whisker plots showing the respective definitive endoderm and hypoblast module scores in each subcluster. Modules were defined as the top 100 most significant markers of the E11.5 definitive endoderm or hypoblast (See Methods). **h**, Violin plot showing the expression of *NODAL* at each time point in the clusters from a. **h**, Predicted *NODAL* cell-cell signalling between

**Extended Data Fig. 11.**
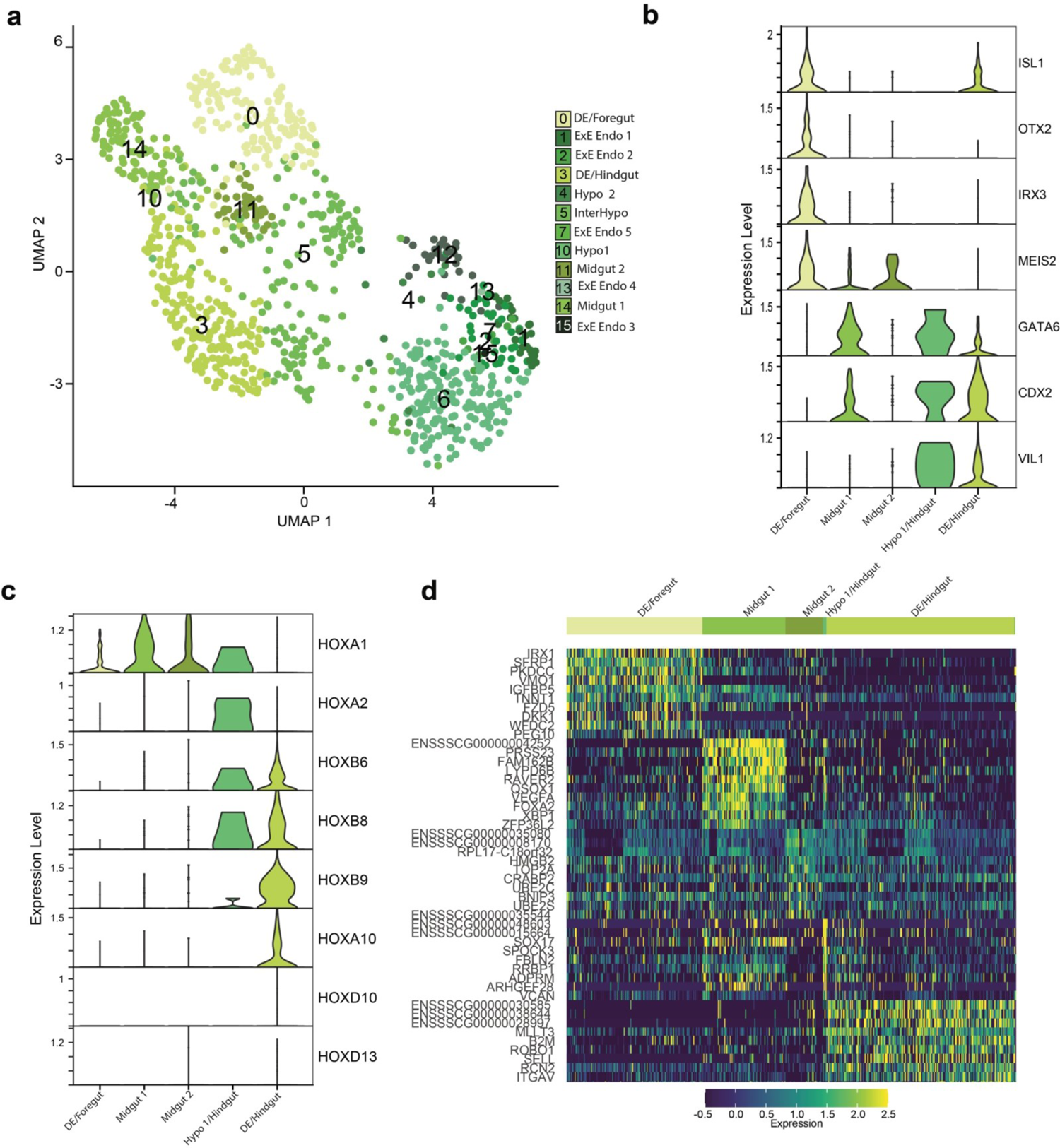
Mature gut types emerge between E14-E15. **a**, UMAP plot showing embryonic and extraembryonic sub-clusters from Fig 10 from E14-E15. **b**, Violin plots of gene expression showing late endodermal subclusters show specific expression of gut markers described by Nowotchin et al^40^. **c**, Violin plots of Hox gene expression showing the distinct anterior-posterior positioning of gut subclusters. **d**, Heat map illustrating the scaled mean expression of the top 10 most significant markers of each gut subcluster.

**Extended Data Fig. 12.**
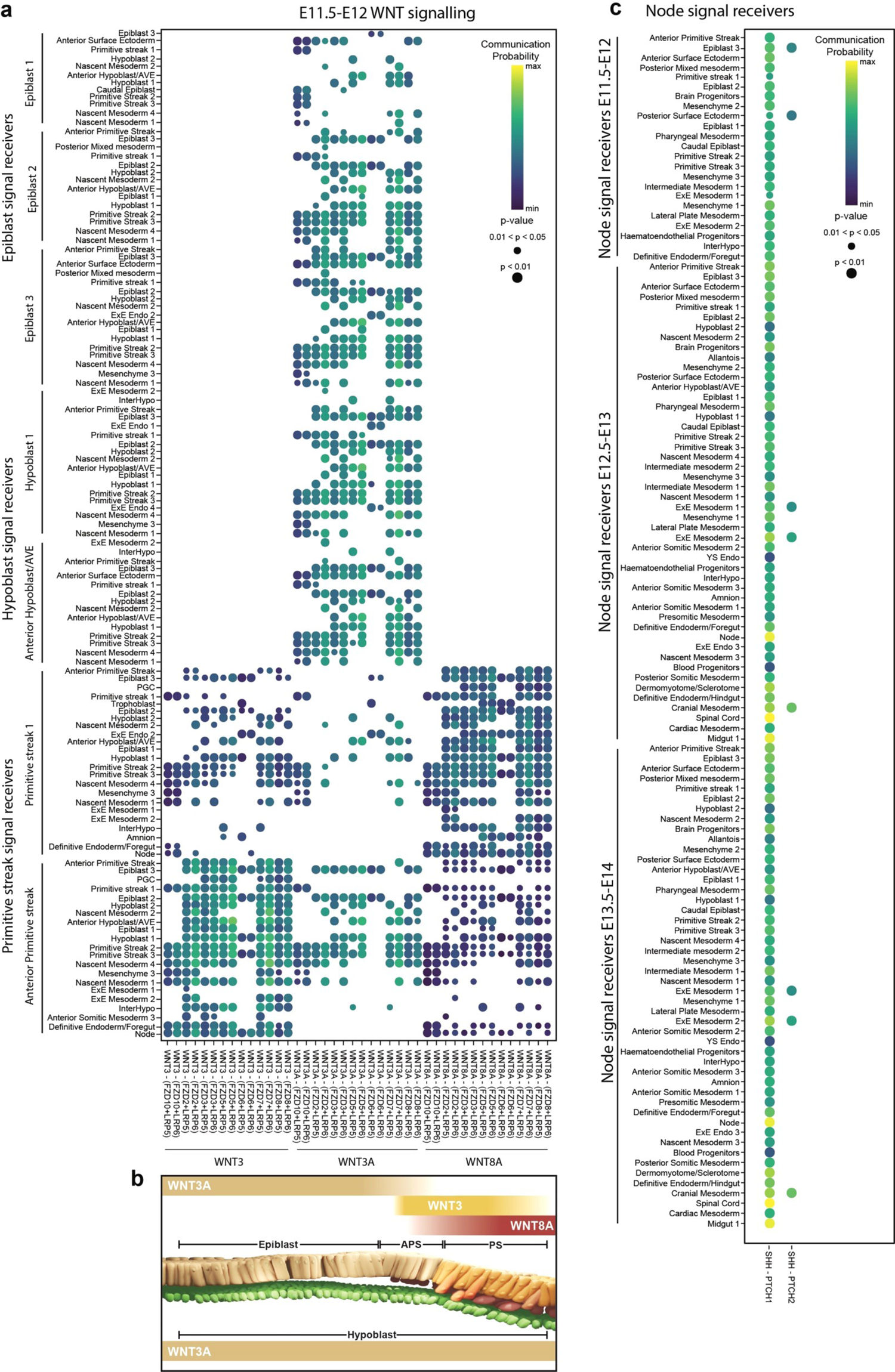
WNT and Nodal signaling in early pig gastrula embryos. **a**, Predicted canonical WNT3, WNT3A and WNT8A cell-cell signalling in E11.5-12 Epiblast, Hypoblast and primitive streak clusters. **b**, Schematic summarising a. **c**, Predicted Hedgehog cell-cell signalling between selected cell types in E11.5-14 embryos.

## Notes

### Competing Interest Statement

The authors have declared no competing interest.

## References

1. Arias, A.M., Marikawa, Y. & Moris, N. Gastruloids: Pluripotent stem cell models of mammalian gastrulation and embryo engineering. Dev Biol 488, 35–46 (2022).

2. Pijuan-Sala, B. et al. A single-cell molecular map of mouse gastrulation and early organogenesis. Nature 566, 490–495 (2019).

3. Mayshar, Y. et al. Time-aligned hourglass gastrulation models in rabbit and mouse. Cell (2023).

4. Ton, M.N. et al. An atlas of rabbit development as a model for single-cell comparative genomics. Nat Cell Biol (2023).

5. Nakamura, T. et al. A developmental coordinate of pluripotency among mice, monkeys and humans. Nature 537, 57–62 (2016).

6. Zhai, J. et al. Primate gastrulation and early organogenesis at single-cell resolution. Nature 612, 732–738 (2022).

7. Alberio, R., Kobayashi, T. & Surani, M.A. Conserved features of non-primate bilaminar disc embryos and the germline. Stem Cell Reports 16, 1078–1092 (2021).

8. Das, S. et al. Generation of human endothelium in pig embryos deficient in ETV2. Nat Biotechnol 38, 297–302 (2020).

9. Telugu, B.P., Park, K.E. & Park, C.H. Genome editing and genetic engineering in livestock for advancing agricultural and biomedical applications. Mamm Genome 28, 338–347 (2017).

10. Sykes, M. & Sachs, D.H. Transplanting organs from pigs to humans. Sci Immunol 4 (2019).

11. Peter, I.S. & Davidson, E.H. The endoderm gene regulatory network in sea urchin embryos up to mid-blastula stage. Dev Biol 340, 188–199 (2010).

12. Sulston, J.E., Schierenberg, E., White, J.G. & Thomson, J.N. The embryonic cell lineage of the nematode Caenorhabditis elegans. Dev Biol 100, 64–119 (1983).

13. Warga, R.M. & Nusslein-Volhard, C. Origin and development of the zebrafish endoderm. Development 126, 827–838 (1999).

14. Hatada, Y. & Stern, C.D. A fate map of the epiblast of the early chick embryo. Development 120, 2879–2889 (1994).

15. Scheibner, K. et al. Epithelial cell plasticity drives endoderm formation during gastrulation. Nat Cell Biol 23, 692–703 (2021).

16. Probst, S. et al. Spatiotemporal sequence of mesoderm and endoderm lineage segregation during mouse gastrulation. Development 148 (2021).

17. Kubo, A. et al. Development of definitive endoderm from embryonic stem cells in culture. Development 131, 1651–1662 (2004).

18. Yasunaga, M. et al. Induction and monitoring of definitive and visceral endoderm differentiation of mouse ES cells. Nat Biotechnol 23, 1542–1550 (2005).

19. D’Amour, K.A. et al. Efficient differentiation of human embryonic stem cells to definitive endoderm. Nat Biotechnol 23, 1534–1541 (2005).

20. Tada, S. et al. Characterization of mesendoderm: a diverging point of the definitive endoderm and mesoderm in embryonic stem cell differentiation culture. Development 132, 4363–4374 (2005).

21. Kinoshita, M. et al. Pluripotent stem cells related to embryonic disc exhibit common self-renewal requirements in diverse livestock species. Development 148 (2021).

22. Thomson, J.A. et al. Embryonic stem cell lines derived from human blastocysts. Science 282, 1145–1147 (1998).

23. Pham, T.X.A. et al. Modeling human extraembryonic mesoderm cells using naive pluripotent stem cells. Cell Stem Cell 29, 1346–1365 e1310 (2022).

24. Luckett, W.P. Origin and differentiation of the yolk sac and extraembryonic mesoderm in presomite human and rhesus monkey embryos. Am J Anat 152, 59–97 (1978).

25. Flechon, J.E., Degrouard, J. & Flechon, B. Gastrulation events in the prestreak pig embryo: ultrastructure and cell markers. Genesis 38, 13–25 (2004).

26. Yang, R. et al. Amnion signals are essential for mesoderm formation in primates. Nat Commun 12, 5126 (2021).

27. Pfeffer, P.L. Alternative mammalian strategies leading towards gastrulation: losing polar trophoblast (Rauber’s layer) or gaining an epiblast cavity. Philos Trans R Soc Lond B Biol Sci 377, 20210254 (2022).

28. Qiu, C. et al. Systematic reconstruction of cellular trajectories across mouse embryogenesis. Nat Genet 54, 328–341 (2022).

29. Ton, M.-L.N. et al. Rabbit Development as a Model for Single Cell Comparative Genomics. *bioRxiv*, 2022.2010.2006.510971 (2022).

30. Tyser, R.C.V. et al. Single-cell transcriptomic characterization of a gastrulating human embryo. Nature 600, 285–289 (2021).

31. Mittnenzweig, M. et al. A single-embryo, single-cell time-resolved model for mouse gastrulation. Cell 184, 2825–2842 e2822 (2021).

32. van de Pavert, S.A. et al. Comparison of anterior-posterior development in the porcine versus chicken embryo, using goosecoid expression as a marker. Reprod Fertil Dev 13, 177–185 (2001).

33. Dobreva, M.P. et al. Amniotic ectoderm expansion in mouse occurs via distinct modes and requires SMAD5-mediated signalling. Development 145 (2018).

34. Hashmi, A. et al. Cell-state transitions and collective cell movement generate an endoderm-like region in gastruloids. Elife 11 (2022).

35. Chapman, D.L. & Papaioannou, V.E. Three neural tubes in mouse embryos with mutations in the T-box gene Tbx6. Nature 391, 695–697 (1998).

36. Chu, L.F. et al. An In Vitro Human Segmentation Clock Model Derived from Embryonic Stem Cells. Cell Rep 28, 2247–2255 e2245 (2019).

37. Guibentif, C. et al. Diverse Routes toward Early Somites in the Mouse Embryo. Dev Cell 56, 141–153 e146 (2021).

38. Kobayashi, T. et al. Principles of early human development and germ cell program from conserved model systems. Nature 546, 416–420 (2017).

39. Rothova, M.M. et al. Identification of the central intermediate in the extra-embryonic to embryonic endoderm transition through single-cell transcriptomics. Nat Cell Biol 24, 833–844 (2022).

40. Nowotschin, S. et al. The emergent landscape of the mouse gut endoderm at single-cell resolution. Nature 569, 361–367 (2019).

41. Kwon, G.S., Viotti, M. & Hadjantonakis, A.K. The endoderm of the mouse embryo arises by dynamic widespread intercalation of embryonic and extraembryonic lineages. Dev Cell 15, 509–520 (2008).

42. Yoshida, M. et al. Conserved and divergent expression patterns of markers of axial development in eutherian mammals. Dev Dyn 245, 67–86 (2016).

43. Rito, T., Libby, A.R.G., Demuth, M. & Briscoe, J. Notochord and axial progenitor generation by timely BMP and NODAL inhibition during vertebrate trunk formation. *bioRxiv*, 2023.2002.2027.530267 (2023).

44. Garcia, M.R. et al. In vitro modelling of anterior primitive streak patterning with hESC reveals the dynamic of WNT and NODAL signalling required to specify notochord progenitors. bioRxiv, 2023.2006.2001.543323 (2023).

45. Martinez Arias, A. & Steventon, B. On the nature and function of organizers. Development 145 (2018).

46. Liu, P. et al. Requirement for Wnt3 in vertebrate axis formation. Nat Genet 22, 361–365 (1999).

47. Martyn, I., Kanno, T.Y., Ruzo, A., Siggia, E.D. & Brivanlou, A.H. Self-organization of a human organizer by combined Wnt and Nodal signalling. Nature 558, 132–135 (2018).

48. Martyn, I., Brivanlou, A.H. & Siggia, E.D. A wave of WNT signaling balanced by secreted inhibitors controls primitive streak formation in micropattern colonies of human embryonic stem cells. Development 146 (2019).

49. Takada, S. et al. Wnt-3a regulates somite and tailbud formation in the mouse embryo. Genes Dev 8, 174–189 (1994).

50. Jin, S. et al. Inference and analysis of cell-cell communication using CellChat. Nat Commun 12, 1088 (2021).

51. McMahon, J.A. et al. Noggin-mediated antagonism of BMP signaling is required for growth and patterning of the neural tube and somite. Genes Dev 12, 1438–1452 (1998).

52. Boulet, A.M. & Capecchi, M.R. Signaling by FGF4 and FGF8 is required for axial elongation of the mouse embryo. Dev Biol 371, 235–245 (2012).

53. Anderson, R.M., Lawrence, A.R., Stottmann, R.W., Bachiller, D. & Klingensmith, J. Chordin and noggin promote organizing centers of forebrain development in the mouse. Development 129, 4975–4987 (2002).

54. Guo, G. et al. Naive Pluripotent Stem Cells Derived Directly from Isolated Cells of the Human Inner Cell Mass. Stem Cell Reports 6, 437–446 (2016).

55. Bergmann, S. et al. Spatial profiling of early primate gastrulation in utero. Nature 609, 136–143 (2022).

56. Satija, R., Farrell, J.A., Gennert, D., Schier, A.F. & Regev, A. Spatial reconstruction of single-cell gene expression data. Nat Biotechnol 33, 495–502 (2015).

57. McGinnis, C.S., Murrow, L.M. & Gartner, Z.J. DoubletFinder: Doublet Detection in Single-Cell RNA Sequencing Data Using Artificial Nearest Neighbors. Cell Syst 8, 329–337 e324 (2019).

58. Trapnell, C. et al. The dynamics and regulators of cell fate decisions are revealed by pseudotemporal ordering of single cells. Nat Biotechnol 32, 381–386 (2014).

59. Van de Sande, B. et al. A scalable SCENIC workflow for single-cell gene regulatory network analysis. Nat Protoc 15, 2247–2276 (2020).

