## Supplementary Figures for "A single-cell atlas of pig gastrulation as a resource for comparative embryology"

Supplementary Fig. 1

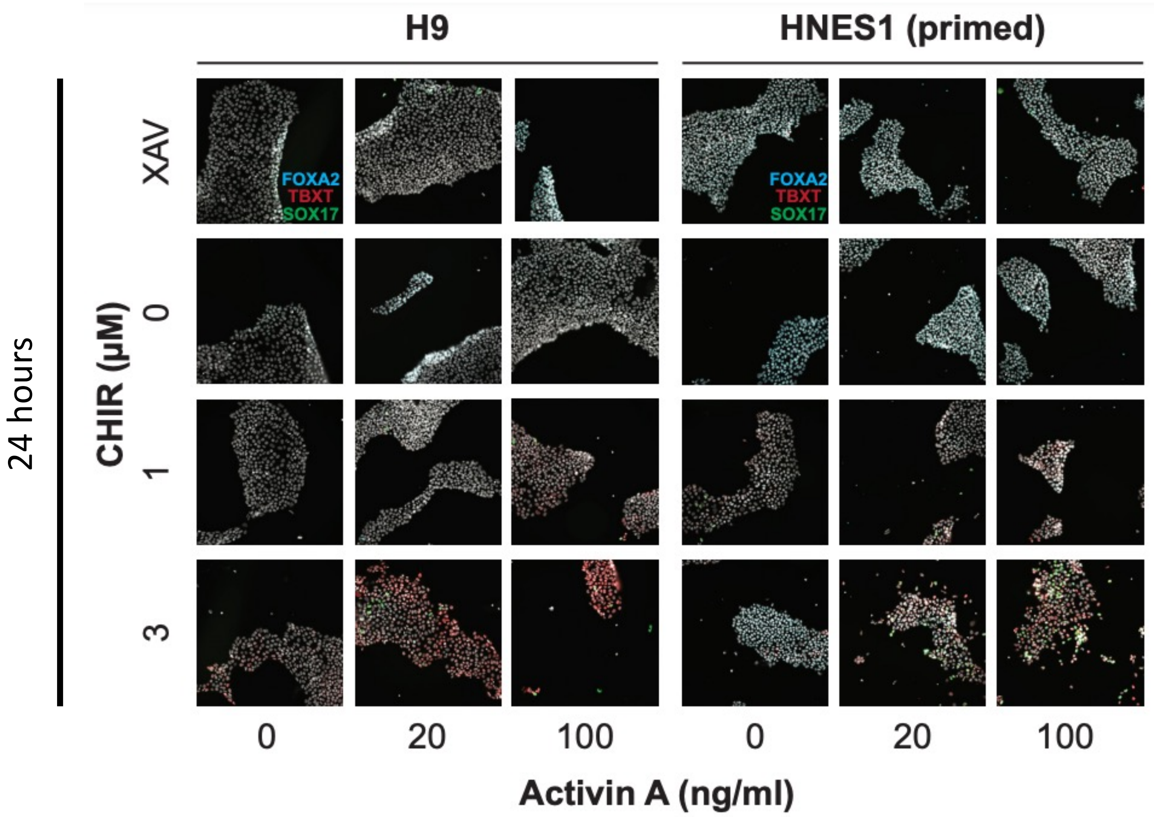


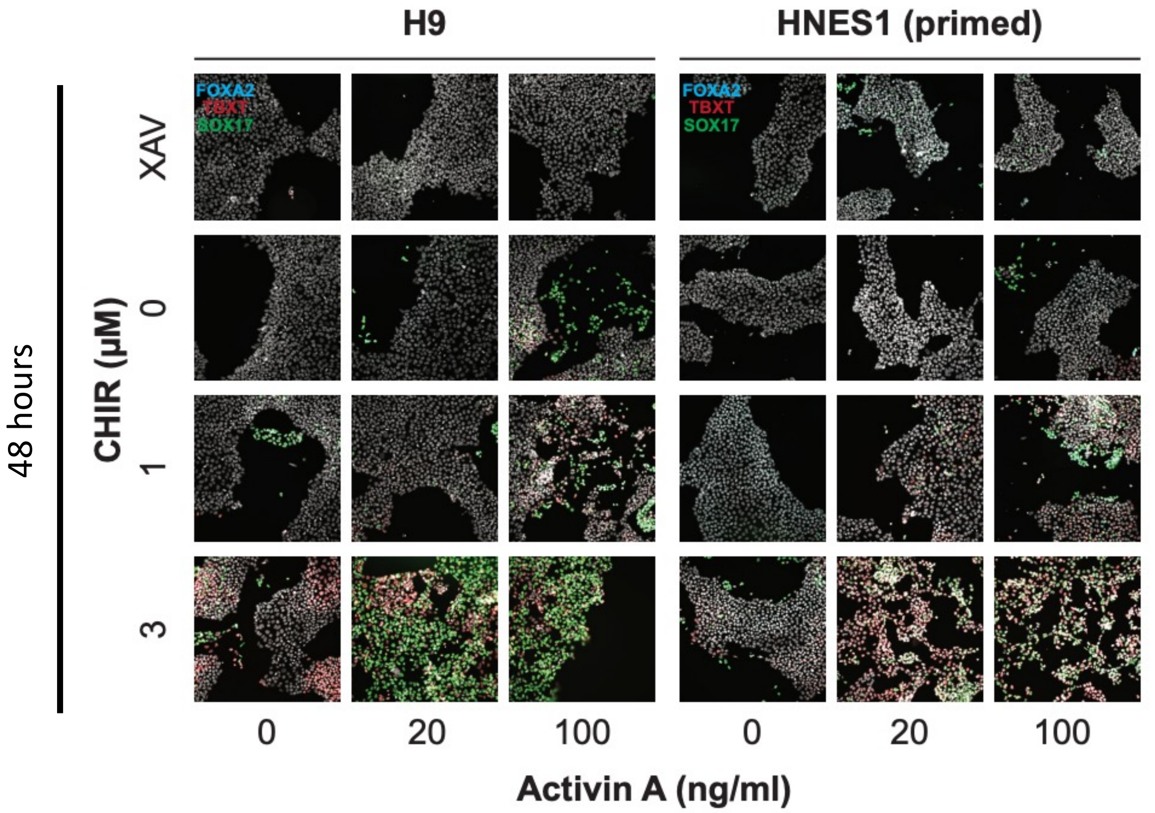


**Supplementary Fig. 1. Co-expression of mesodermal and endoderm markers.** Representative images of H9 and HNES1 hESC lines 24 (Top) and 48 hours (Bottom) after treatments with differing amounts of XAV, CHIR and/or Activin A. Cells were then stained for FOXA2, TBXT and SOX17 expression.

Supplementary Fig.2

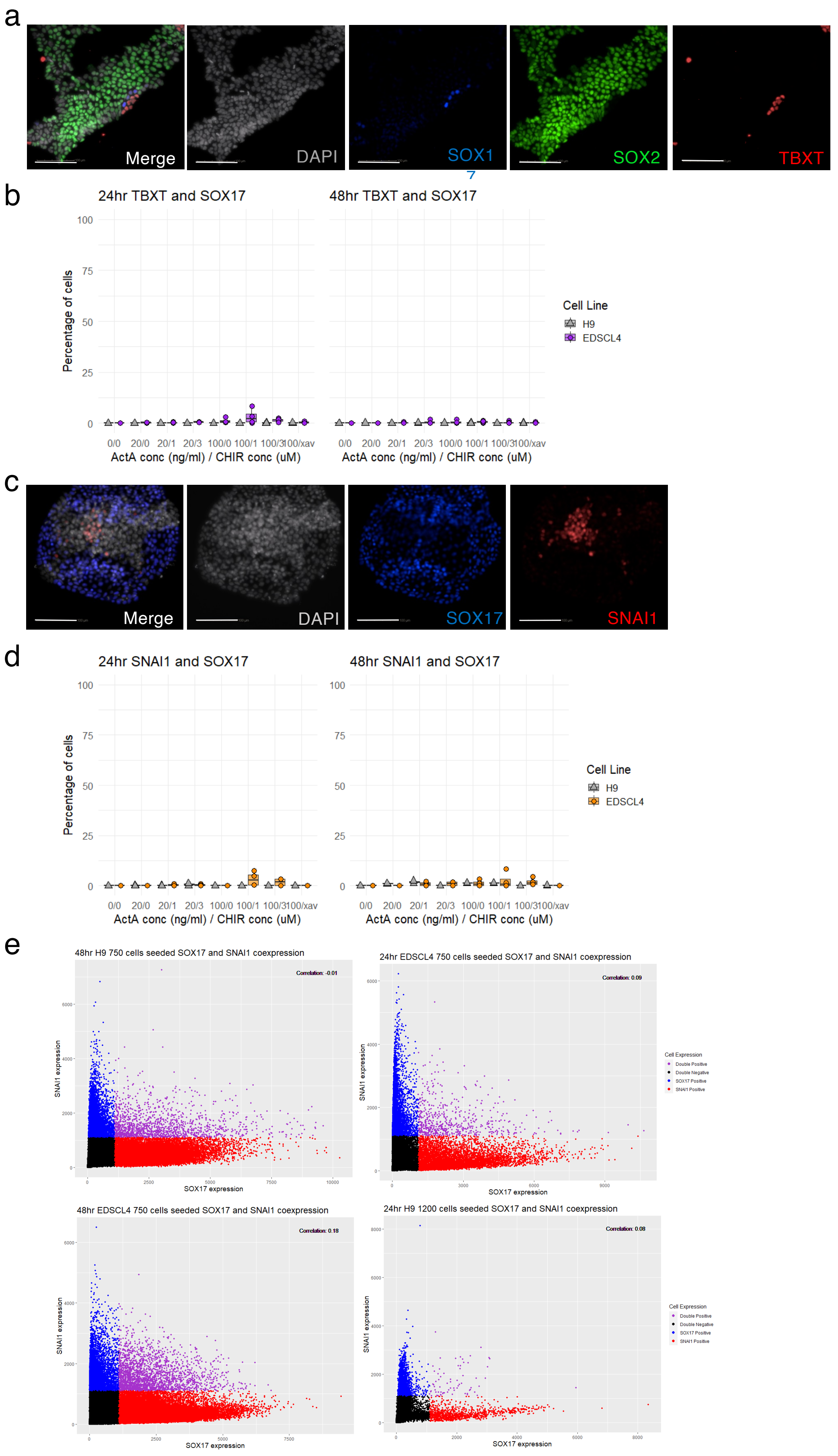


**Supplementary Fig. 2. Co-expression of endoderm markers. a,** representative images of EDSCL4 after 8 hours of induction with ActA (100ng/ml) and CHIR (1$\mu$M). Scale bar represents 100$\mu$m. **b**, Quantification of SOX17 and TBXT co-expression across the 2D experiments. **c,** Representative images of EDSCL4 after 48 hours of induction with ActA (100ng/ml) and CHIR (1$\mu$M). Scale bar represents 100$\mu$m. **d**, Quantification of SOX17 and SNAI1 co-expression across the 2D work. **e,** Selection of correlation graphs between SNAI1 and SOX17.Correlation value calculated via Pearson's correlation.
